# MicroRNAs are deeply linked to the emergence of the complex octopus brain

**DOI:** 10.1101/2022.02.15.480520

**Authors:** Grygoriy Zolotarov, Bastian Fromm, Ivano Legnini, Salah Ayoub, Gianluca Polese, Valeria Maselli, Peter J. Chabot, Jakob Vinther, Ruth Styfhals, Eve Seuntjens, Anna Di Cosmo, Kevin J. Peterson, Nikolaus Rajewsky

## Abstract

Soft-bodied cephalopods such as the octopus are exceptionally intelligent invertebrates with a highly complex nervous system that evolved independently from vertebrates. Because of elevated RNA editing in their nervous tissues, we hypothesized that RNA regulation may play a major role in the cognitive success of this group. We thus profiled mRNAs and small RNAs in 18 tissues of the common octopus. We show that the major RNA innovation of soft-bodied cephalopods is a massive expansion of the miRNA gene repertoire. These novel miRNAs were primarily expressed in neuronal tissues, during development, and had conserved and thus likely functional target sites. The only comparable miRNA expansions happened, strikingly, in vertebrates. Thus, we propose that miRNAs are intimately linked to the evolution of complex animal brains.

**One-Sentence Summary:** miRNAs are deeply linked to the emergence of complex brains.

## Main Text

Coleoid (soft-bodied) cephalopods (octopuses, squids, and cuttlefishes) possess elaborate nervous systems both in terms of size and organization (*1*–*4*). Understanding the molecular mechanisms behind the evolution of the coleoid nervous system thus offers the opportunity to discover general molecular design principles behind morphological and behavioral complexity in animals. Octopus (*5*) and squid (*6*) genomes do not show signs of whole-genome duplications, and the intronic architecture, as well as protein-coding content, were found to largely resemble those of other related invertebrates (*7*). Recently, it was shown that coleoids extensively use A-to-I RNA editing (*8, 9*) mediated by ADAR enzymes (“adenosine deaminases acting on RNAs”) (*10*) to re-code their neuronal transcriptomes. Because extensive editing is not abundant in other mollusks including *Nautilus*, a cephalopod and the living sister group of the coleoids with a simpler nervous system, this process has been hypothesized to drive the cognitive success of coleoids (*9*), perhaps by providing a mechanism to expand and regulate the coding repertoire of mRNAs. However, it is difficult to explain the evolution of complex heritable traits by the actions of a single trans-acting factor, and indeed it has been proposed that the editing phenomena in coleoids are mainly non-adaptive ((*11*), but see (*12*)). Because ADARs interact and regulate many classes of RNAs (for example, the silencing of transposon RNA (*13*), the biogenesis of circular RNAs (circRNAs) (*14*), and defense against viral RNAs (*15*), we hypothesized that post-transcriptional regulation of RNA in general is potentially linked to the evolution of the complex nervous system of the coleoid cephalopods.

Thus, we systematically quantified major modes of post-transcriptional regulation across 18 tissues of adult octopus (Fig 1A, B, Table S1 and S2). For each mode of regulation, we also checked if A-to-I editing adds complexity to regulation. Briefly (see our Supplementary Text for an in-depth presentation), we combined mRNA shotgun and two full-length mRNA sequencing methods (Iso-seq from PacBio and FLAM-seq (*16*)) to produce a high-quality dataset of 56,579 mRNA isoforms covering 10,957 reference genes (Supplementary Data 1). Both in neuronal and non-neuronal tissues, the majority of A-to-I editing occurred in the introns and 3’-untranslated regions (3’-UTRs) of mRNAs, consistent with the elevated presence of ADAR substrates (hairpin structures) in these regions compared to coding sequences (Fig. S1). We found that alternative splicing was highest in neural tissues, as expected, and that A-to-I editing very rarely altered splice sites (Fig S2, Table S3). CircRNAs, were expressed at overall low levels, consistent with the reported repression of circRNA biogenesis by ADAR (*14, 17*). When analyzing poly-A tails with FLAM-seq, we discovered that poly-A tails from the octopus testes were significantly shorter than in any other tissue and, surprisingly, contained a high fraction of guanosines, a phenomenon not seen in other species (Supp. Text and Fig. S3). 3’-UTRs had a median length of 380 nt -longer than in well-studied invertebrate model systems.

**Fig. 1.**
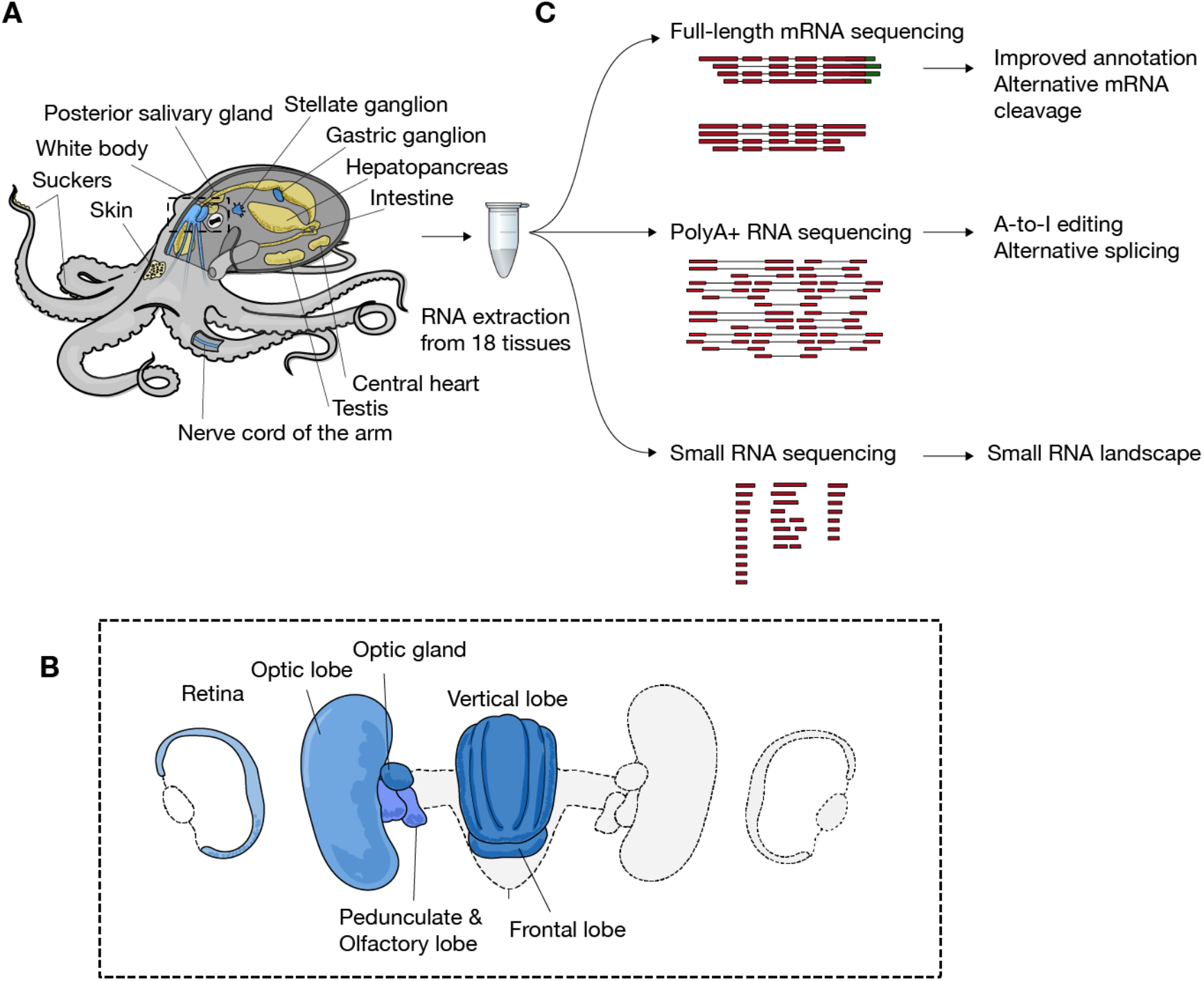
RNA profiling of a common octopus. (**A**) Schematic representation of tissues sampled in the study. Neuronal and non-neuronal tissues are colored in blue and yellow, respectively. Inset (**B**): brain and surrounding structures. (**C**) Main sequencing methods and computational analyses used in this study.

In summary, the transcriptome of a common octopus does not show major departures from other invertebrates in terms of alternative splicing diversity and rates, as well as in mRNA cleavage and polyadenylation. The most outstanding feature was 3’-UTR length and we thus turned our focus on miRNAs which are known to bind 3’-UTRs and to have complex patterns over evolution history (*18*).

### A massive expansion of the miRNA repertoire in coleoid cephalopods

When annotating miRNAs from small RNA sequencing data, a fundamental problem is the detection of a large number of lowly expressed small RNAs that are probably background products of the miRNA biogenesis pathway without functional importance (*19*). To focus on miRNAs that are functionally important, we removed all *vulgaris* small RNAs which we could not find in a whole-body small RNA sequencing dataset of the octopus *O. bimaculoides*, which split from *O. vulgaris* ∼50 million years ago (*20*) (Methods). We thus identified a total of 177 conserved octopus miRNAs. We stress that this is likely an underestimate of the number of functional octopus miRNAs as our sequencing data are incomplete and we are missing functional miRNAs that may have evolved during the past 50 million years in one or the other octopus species. However, we recovered 46/48 miRNA families expected to be present in octopus given its phylogenetic position (Methods). Two families (miR-1989 and miR-242) were not found in the expression data or genomes from both octopus species, and neither were present in either the genome (*6*) or in the miRNA-seq of a bobtail squid *E. scolopes*, but were present in the *Nautilus* genome (*21*). These two miRNA loci were thus likely lost in the coleoid lineage. In total, 39% (69/177) of predicted miRNA genes could be assigned to known miRNA families described in other animals. Out of 108 potentially novel miRNAs, 12 were found in the genome of *Nautilus* and a squid, and thus represent the cephalopod miRNA set (Fig 2A). An additional 51 novel miRNA genes (grouped into 42 miRNA families) are shared between the octopuses and squid, and thus represent miRNAs that emerged in coleoid lineage. Finally, the remaining 42 miRNA genes (35 families) are restricted to the *Octopus* lineage.

**Fig. 2.**
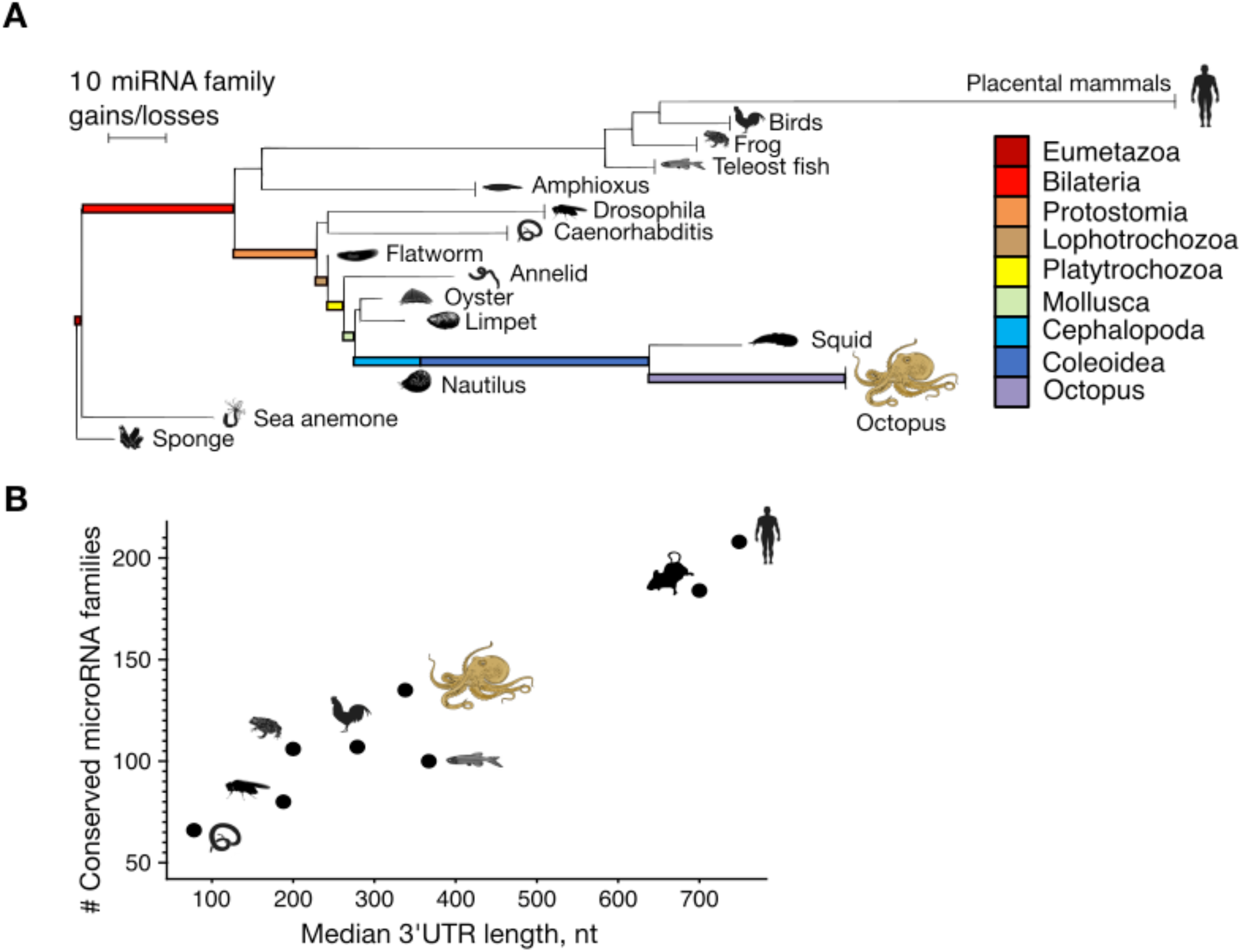
Massive expansion of the miRNA repertoire in cephalopods. (**A**) Phylogeny of several animal groups with the branch lengths between nodes, or from a node to an extant species, reflecting the gains of miRNA families minus the losses (Methods). Vertical lines at the end of the branches indicate the shared complement of the indicated taxon as deposited in MirGeneDB (*46*); the other branches lead to single species (sponge: *A. queenslandica*; sea anemone: *N. vectensis*; flatworm: *S. mediterranea*; annelid: *C. teleta*; oyster: *C. gigas*; limpet: *L. gigantea*). (**B**) Number of miRNA families (excluding species-specific novel families) versus median 3’UTR length in selected animals. For instance, “Human” represents the number of miRNA families annotated in genus *Homo*. Median lengths of 3’UTRs were computed from genome annotations (Methods).

This dramatic expansion of the miRNA gene repertoire in soft-bodied cephalopods (at least 89 gene families) is the largest gain of shared miRNA families known within the invertebrates, and the total number of miRNA families in the octopus genome (135) is on par with that found in vertebrates (minus placental mammals) including chicken (107 families), African clawed frog (106), or zebrafish (100) (Fig 2A). An evolutionary expansion of the number of miRNAs is generally linked to an expansion in the length of 3’-UTRs (*22*), the targets of miRNAs. Indeed, when using our measured 3’-UTR lengths in octopus and graphing the number of conserved miRNAs versus median 3’-UTR length for octopus and other species, the octopus data fit nicely into the expected position (Fig 2B).

### Novel miRNAs are specifically expressed in neural tissues and during development

We next investigated tissue expression patterns of octopus miRNAs as a function of their evolutionary age. Deeply conserved bilaterian miRNAs recapitulated known tissue expression patterns (*23*) (Table S5). The majority of cephalopod and coleoid-specific miRNAs was expressed, as expected, at overall lower levels than older miRNAs (Fig. S4) (*24, 25*). However, novel miRNAs were primarily expressed in the nervous tissues of the animal (Fig. 3). Of the 51 miRNAs of coleoid origin, 45 have their maximum of expression in one or more neural tissues. In these tissues, they are expressed, on average, at 13 times higher levels than in non-neuronal tissues (Fig. S4 B and C, Fig. S5). In fact, the sampled non-neuronal tissue with the highest coverage (“Suckers tip”, 50 M reads; 53.6%) had a lower proportion of captured novel microRNAs than the neuronal tissue with the lowest coverage (“Pedunculate and Olfactory lobe”, 30 M reads; 57%). Thus, the fact that the novel miRNAs are most specifically expressed in neural tissues is not due to a potential tissue sampling bias (Suppl. Text).

**Fig 3.**
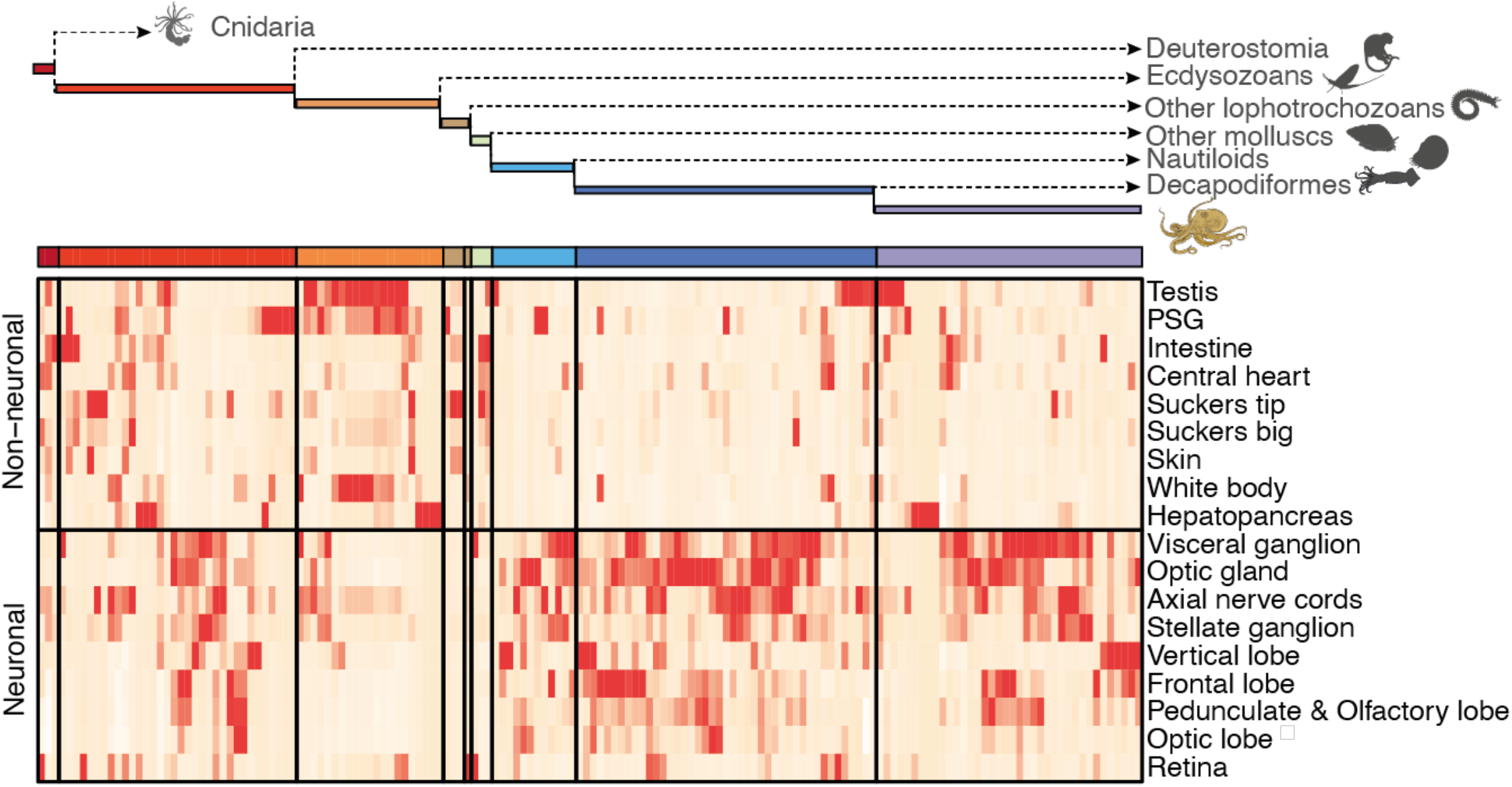
Novel, conserved octopus miRNAs are specifically expressed in neuronal tissues. A simplified phylogenetic tree showing the number of miRNAs that evolved from the time bilaterians split from cnidarians to the last common ancestor of the two considered *Octopus* species. Color code as in Fig. 2. For each miRNA (columns), its expression distribution across tissues (rows) in both neural and non-neural tissues and the corresponding Z-scores were computed. Columns within each bin were hierarchically clustered based on the Z-scores (extended version: Fig. S3A).

If these novel miRNAs contribute to the evolution of the octopus brain, they would be expected to be expressed during neural development. To test this prediction, we profiled small RNA expression at the late stages of *O. vulgaris* development before hatching. Moreover, immediately after hatching, we sequenced small RNAs from whole-body hatchlings as well as isolated brains (Fig. 4). Novel coleoid miRNAs were robustly expressed during development and had the largest contribution (compared to older miRNAs) in the hatchling’s brains. Strikingly, this tissue had the highest relative proportion (∼70%) of a miRNA transcriptome devoted to evolutionary novel miRNAs of all 22 tissues sequenced in this study (Fig. S6). Together, our data suggest that indeed novel coleoid miRNAs contribute to the development of the octopus brain.

**Fig. 4.**
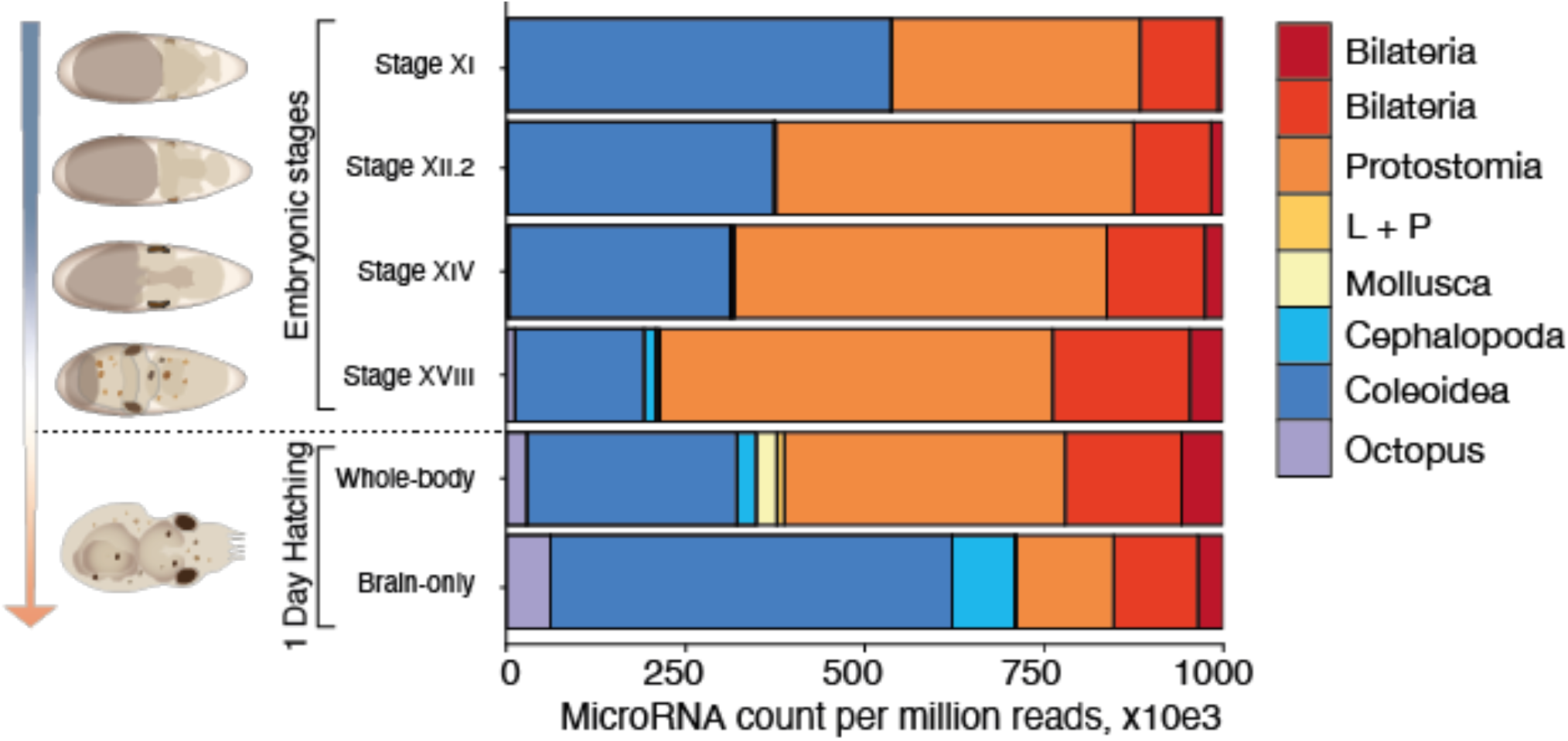
Novel, conserved octopus miRNAs are highly expressed during development with peak expression in brain. miRNA expression (in sequencing reads per million) colored by the phylogenetic node of origin. Samples were obtained by developmental stage of *O. vulgaris* (*47*). These samples cover the organogenic stages of *O. vulgaris* development (Stage XI -Stage XVIII) when most of the embryonic growth occurs, as well as the whole-body and brain of one-day-old paralarvae when the growth of the larval brain commences. An extended version of this figure with per-library depth and numbers of detected miRNAs is available in Fig. S5. “L + P” refers to the collective miRNAs that evolved in lophotrochzoans and platytrochozoans (see Fig. 2).

### Target sites of novel miRNAs are conserved

If miRNA target sites are conserved across sufficiently large evolutionary distances, it is likely that these sites are functionally important. Thus, to show that the shared miRNA complement of the two *Octopus* species are functional, we asked if their target sites are conserved between these two species. To this end, we defined “miRNA response elements” (MREs) as an octamer starting with adenosine followed by a heptamer Watson-Crick complementary to position 2-8 of the miRNA (Fig. 5A) (*26, 27*). These MREs generally mediate the strongest regulatory effect when bound by the respective miRNA. Indeed, predicted MREs shared between the two octopus species showed higher conservation rates compared to the control 8-mers (Fig. 5B, Methods, Supp. Data 2). As expected (*28*), this signal disappeared when miRNA:target pairs were not co-expressed (Fig. 5B, Methods, Supp. Data 2) strongly suggesting that the conservation of MRE’s is indeed caused by the functional interaction between the miRNA and the MRE in the respective tissues. Finally, MREs of phylogenetically younger miRNA families were, on average, less conserved than MREs from older miRNA families (i.e. miRNAs of protostome or bilaterian origin) (Fig. 5C), consistent with their generally lower expression levels and potentially lower selection pressure to maintain their target sites (*29*). Overall, we conclude that the novel octopus miRNAs are functional and exert function, at least in part, by canonical seed-pairing mechanism.

**Fig. 5.**
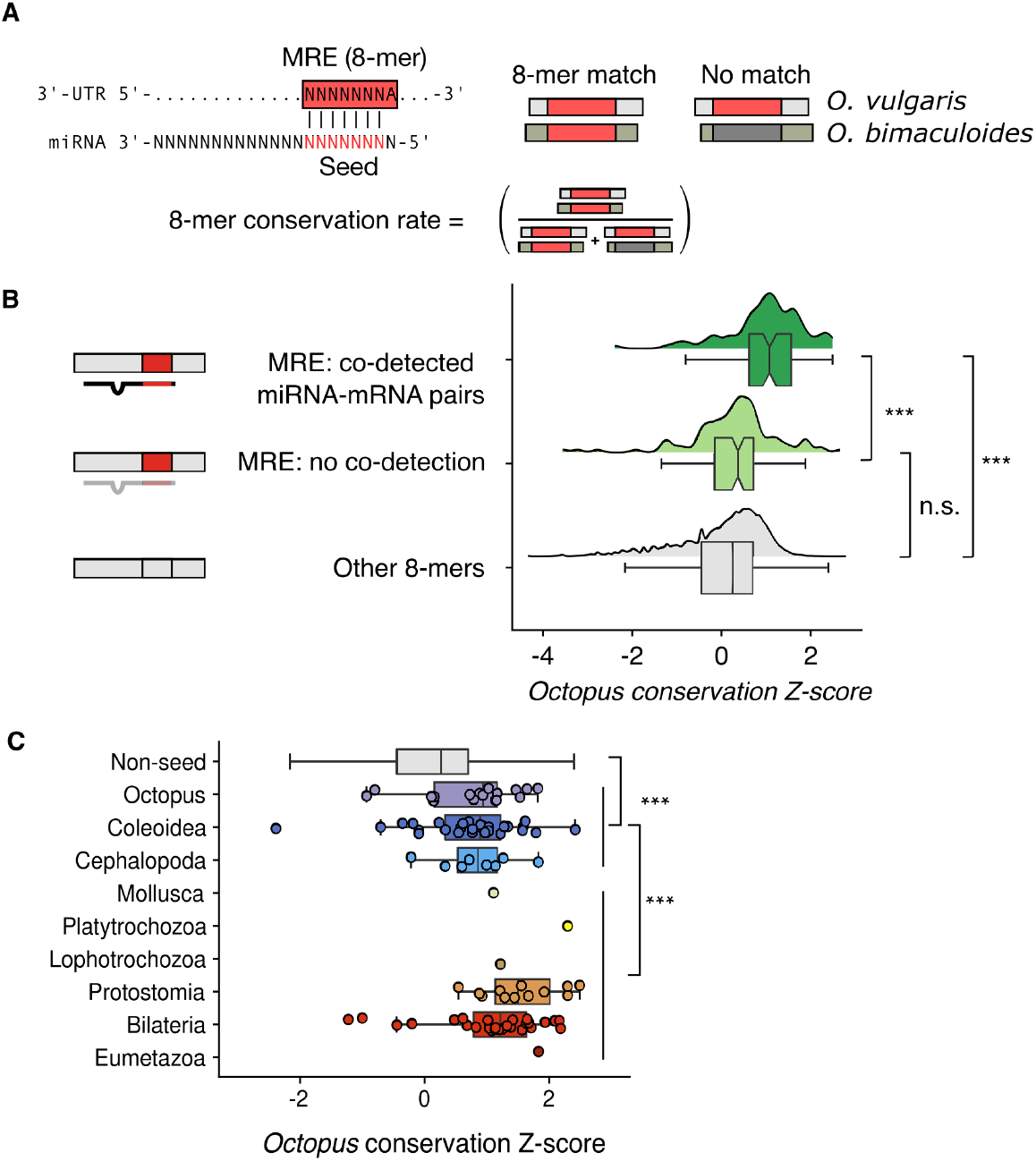
Target sites of novel miRNAs are conserved and co-expressed with the respective miRNA. **(A)** Definition of “miRNA response elements” (or “8-mer”) and their evolutionary conservation. The 8-mer conservation rate is defined as the percentage of occurrences in 3’ UTRs, where a particular 8-mer (red) is matched by exactly the same 8-mer at the same position in the aligned orthologous 3’-UTR. **(B)** Shown here, for novel octopus miRNAs (conserved between *vulgaris* and *bimaculoides*), is the MRE conservation rate in units of a standard Z score. Co-expression is defined as a mRNA with a MRE and the respective miRNA co-detected in at least one tissue at 10 and 100 counts per million, respectively (Methods). Co-expressed miRNA-MRE pairs are statistically more highly (p < 0.001) conserved than non-co-expressed pairs or control 8mers which were not related to any MRE in the octopus. **(C)** As expected, MREs conservation rates are higher for evolutionarily older miRNA families. In (B) and (C), statistical significances: Mann-Whitney U test with Bonferroni correction for multiple hypothesis testing (n.s.: p > 0.05, ***: p < 0.001).

### In octopus, A-to-I editing is decoupled from miRNA function

We asked if A-to-I editing is potentially modulating miRNA function in the octopus. This could occur by (i) editing the miRNAs themselves and/or (ii) editing miRNA target sites in 3’-UTRs (destroying or creating them). Briefly, we found no evidence for any functionally important editing of miRNAs (Suppl. Text). We could detect only 5 miRNAs with an estimated A-to-I editing frequency in the seed sequence above 1% (but never more than 4.8%) (Fig S7 and Methods). Similarly, we found that A-to-I editing events with the potential to destroy miRNA target sites (MREs) happen rarely. Out of 10,053 MREs conserved between the two octopus species and having sufficient RNA-seq coverage, only 39 (0.3%) harbored editing events (Methods, Suppl. Text). Finally, we found no higher conservation for 8-mers potentially becoming a MRE by A-to-I editing compared to control sequences (Fig. S8, Suppl. Text). This suggests that *de novo* creation of MREs or disruption of existing MREs by editing is, if existent, a rare phenomenon.

## Discussion

Given the generality of its coleoid protein-encoding genomic repertoire including its transcription factor repertoire (Table S2) (*5, 6, 20*), in addition to the reported elevated rates of A-to-I editing in coleoid neural tissues (*8, 9*), we hypothesized that RNA regulation in general might be involved in driving the dramatic increase in the complexity of the coleoid nervous system. Our data and analyses argue that in terms of alternative splicing diversity and rates (including back-splicing that generates circRNAs), as well as mRNA cleavage and polyadenylation patterns, there is no major departure from other invertebrates. Further, we find no evidence for substantial editing in miRNA seed sequences, nor in potential target sites either in the abrogation of a genetically encoded site or in the creation of a newly relevant site (Fig. S7, S8). Of course, A-to-I editing may still be functionally important in individual cases, especially in terms of potential *cis*-regulatory sites for RNA-binding proteins that might be edited in functionally important ways.

On the other hand, a clear distinction in RNA regulation between coleoid cephalopods and all other known invertebrates is reflected in the dramatic expansion of their miRNA repertoire. The conservation of over 50 miRNA loci in both the squid and octopus lineages since they diverged from one another nearly 300 million years ago (*20*) coupled with the 3’-UTR (Fig. 2B), miRNA expression (Figs. 3, 4) and target site (Fig. 5) analyses discussed above, all strongly suggest that these miRNAs are functionally important during the development of the coleoid nervous system. Like in virtually all other increases to a miRNA repertoire, both the source and evolutionary pressures for the rise of these novel miRNA loci is not known; whole genome duplications can be ruled out (*5, 6*), and scenarios may apply where novel miRNAs with functionally beneficial target sites that create new regulatory circuits evolve from weakly expressed precursors by purging of deleterious target sites (*29*). Nonetheless, once under selection, miRNAs in general are believed to improve the robustness of the developmental process (*30*–*34*), increasing the heritability of the interaction (*35*–*37*), which might then allow for the evolution of new cell types (*38*) and ultimately morphological and behavior complexity (*39, 40*). Indeed, with respect to the development of the nervous system, we note that at least in vertebrates, miRNA are known to have highly complex expression patterns with, for example, miRNA transcripts localized to the synapse and modulating their function (*41*). Further, new pathways have been identified that operate in neurons including highly conserved pathways that trigger the destruction of a miRNA bound to a target site of a specific architecture involving extended complementarity beyond the seed site (*42*–*45*). Although it remains to be seen whether these types of pathways operate in coleoids, the striking explosion of the miRNA gene repertoire in coleoid cephalopods may indicate that miRNAs and, perhaps, their specialized neuronal functions, are indeed deeply linked and possibly required for the emergence of complex brains in animals.

## Notes

## Acknowledgments

We thank members of Rajewsky lab, in particular, Cledi Alicia Cerda Jara for her help and support with experiments and Marcel Schilling and Marvin Jens for their support with bioinformatic analyses; Victoria Symkina for her drawing of an octopus. G. Z. thanks the Rajewsky lab for hospitality and his father L. D. Zolotarov for financial and moral support.

## Data availability

Sequencing data associated with this study have been deposited at GEO (GSE192550) and SRA (PRJNA791920). The *O. sinensis* transcriptome annotation generated in this study is available as Supplementary Data 1. MiRNA predictions and their corresponding tissue expression patterns for *O. vulgaris, O. bimaculoides, E. scolopes* and *N. pompilius* are available at MirGeneDB (*46*).

## Author Contributions

G.Z. and N.R. designed this study. G.P., V.M., and A.d. C. provided adult *O. vulgaris* tissue samples; R.S. and E.S. provided *O. vulgaris* embryonic and hatchling RNA samples; G.Z., I.L. and S.A. produced all *O. vulgaris* sequencing data; J.V. and K.P. produced all *O. bimaculoides* and *E. scolopes* sequencing data. B.F., P.C., and K.P. annotated all miRNA genes. G. Z. performed all other data analyses, supervised by N.R. G.Z., K.P., and N.R. wrote the manuscript with input from all authors.

## Funding Information

G. Z. and all sequencing costs for *O. vulgaris* were supported by a DFG Award to N.R. (“Leibniz award”). Costs for the sequencing the *O. bimaculoides* and *E. scolopes* libraries and the development of miRNA annotation tools were supported by grants from the NSF, NASA-Ames and Dartmouth College to K.P. B.F. is supported by the Tromsø forskningsstiftelse (TFS) and was supported by the Strategic Research Area (SFO) program of the Swedish Research Council (VR) through Stockholm University.

## Notes

### Competing Interest Statement

The authors have declared no competing interest.

