## Supplementary Text, Methods and Figures for "MicroRNAs are deeply linked to the emergence of the complex octopus brain"

### Supplementary Text, Supplementary Figures and Methods

#### Supplementary Figures


- Figure S1 - Genomic targets of A-to-I editing
- Figure S2 – Alternative splicing rates across tissues
- Figure S3 - mRNA cleavage and polyadenylation
- Figure S4 - Extended expression patterns and A-to-I editing of miRNAs
- Figure S5 - Sequencing depth and miRNA discovery
- Figure S6 - Fraction of transcriptome dedicated miRNAs of different phylogenetic ages
- Figure S7 – Editing of miRNAs
- Figure S8 - Conservation of 8-mers potentially convertible to MRE by A-to-I editing


#### Supplementary Tables and Data


- Table S1 – RNA sequencing datasets generated in the study
- Table S2 – Transcription factor counts in the genome
- Table S3 - Alternative splicing and A-to-I editing
- Table S4 - Editing levels of miRNA seeds
- Table S5 – Recapitulation of conserved miRNA of expression patterns
- Supplementary Data 1 – Genome annotation
- Supplementary Data 2 – Conservation proportions of MREs
- Supplementary Data 3 – MiRTrace quality control of small RNA-seq libraries

### Supplementary Text

#### RNA quality assessment

Agilent Bioanalyzer has been used to assess the quality of RNA extracted from the tissues. However, the main quality metric of Bioanalyzer, an RNA integrity Number (RIN), could not be used as it has been developed using mouse and human RNA samples where two ribosomal fragments are expected to dominate total RNA profile. As in the majority of other protostomes(*48*, *49*), 28S RNA in cephalopods bears the so-called “hidden break”. Upon denaturation, the 28S rRNA is broken down to two fragments. With an 18S fragment, a total RNA profile of octopus RNA then consists of 3 fragments which makes it impossible for the algorithm to determine RNA quality properly. An example of a profile of intact RNA is shown below (note the disappearance of the 3rd peak upon heat denaturation, the size scale is shifted which is an artifact):

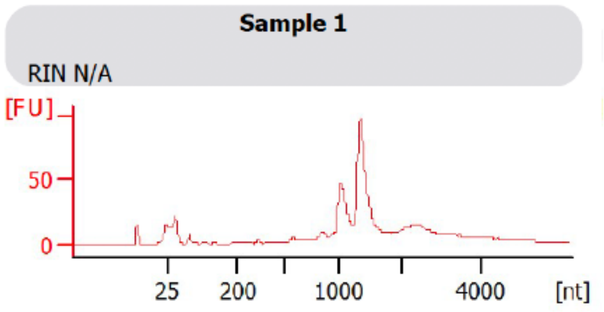


No heat denaturation
hepatopancreas

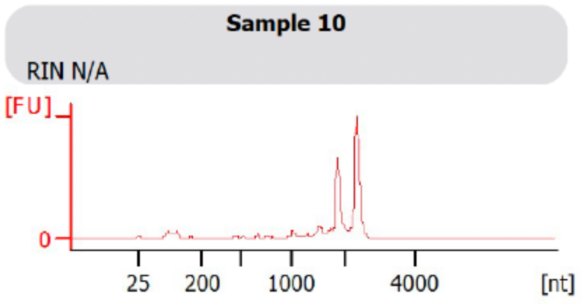


Heat denaturation ( 70C, 3 min )
hepatopancreas

Nevertheless, degraded RNA can be easily distinguished from intact RNA by observing a region between 5S rRNA and the major peaks. In degraded samples, this region will accumulate irregular peaks corresponding to the degradation fragments or larger ribosomal RNAs. An example for a partially and fully degraded samples are show below:

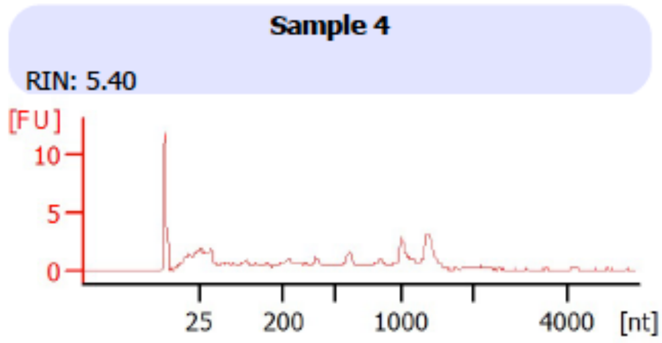


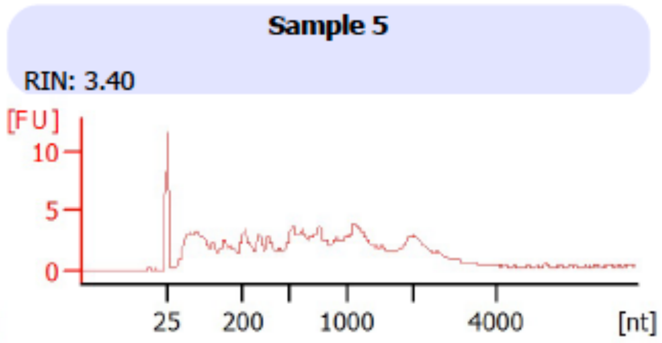


Thus, only samples with clear two or three peaks and no peaks in the intermediate fraction have been selected for the library preparation.

#### Alternative splicing in the octopus genome

Systematic differences in alternative splicing intensity have been reported between metazoans and unicellular holozoans, early-branching non-bilaterian animals, and bilaterians (*50*) as well as within vertebrate lineage (*51*). In particular, exon-skipping events are widespread among metazoans, with the highest rates observed in bilaterian animals and, in particular, vertebrates (*50*). Such increases in the exon-skipping rates have been argued to contribute to an increased phenotypic complexity perhaps by expanding proteome diversity in those lineages (*52*, *53*). An initial assembly and annotation of the *O. bimaculoides* genome suggested no particularly rich alternative splicing of mRNAs. Genes that are highly alternatively spliced in vertebrates have more isoforms in the octopus genome as well, but the number of such isoforms was lower (*5*). In the same study, *O. bimaculoides* has been found to retain 85% of the ancestral introns and the number of novel introns has been found to be comparable to that in other "slow-evolving" spiralians such as *Lottia* and *Capitella*. Here we investigated the extent of alternative splicing by comparing exon skipping and intron retention rates across the tissues of *O. vulgaris* and *O. bimaculoides* (*5*).

Three million sequencing reads generated by two full-length RNA-seq methods (Iso-seq and FLAM-seq (*16*), (*47*)) were used to build a catalog of high-quality mRNAs isoforms (Methods). The isoforms were then filtered (Methods) to produce a final set of 59,579 mRNA isoforms associated with 10,957 reference genes out of 25,335 present in the genome. After filtering, and in combination with a current genome annotation, we estimate that the octopus genome encodes ~80,000 alternative mRNA isoforms. An updated annotation that combines original genome annotation with the isofroms obtained in this study is available as Supplementary Data 1.

We have used our newly assembled transcriptome to characterize alternative splicing in the genome of *O. sinensis* (Methods). *O. sinensis* chromosome-level genome assembly ASM634580v1 (*54*) was used due to its much higher completeness in comparison to a draft genome assembly of *O. vulgaris* (*55*). Both species belong to the same species complex and only recently have been recognized as separate species belonging to the same species complex (*56*, *57*) .

In the obtained annotation, 9769 out of 25,335 protein coding genes (38.6%) encoded multiple mRNA isoforms. As in other bilaterians (*50*), the predominant alternative splicing event in the octopus is exon skipping (6513 cases) followed by alternative transcription start and termination. (4773 and 2580 cases respectively), alternative first exon (2313) and intron retention (1927) (Methods). Individual tissues varied in alternative splicing abundance with neuronal tissues often having higher exon-skipping rates (Methods, Fig. S2). We checked whether A-to-I editing may contribute to an increased number of mRNA isoforms by affecting alternative splicing of messenger RNAs. Splicing requires donor and acceptor splicing signals in the mRNA (GT and AG respectively) as well as other splicing-regulatory sequences within introns (branch point, polypyrimidine tract, etc.) As A-to-I editing is thought to occur co-transcriptionally, it is feasible for splice sites to be created by adenosine deamination. A notable example of such event in the autoregulatory loop of ADAR2 itself, where adenosine deamination in ADAR2 mRNA creates a proximal splicing acceptor site and results in a premature translation termination due to a frame shift thus possibly acting in self-inhibitory fashion (*58*). However, in human cell lines, ADAR enzymes have been found to target splicing-related motifs only rarely (*59*). If splice donor or/and acceptor site are created by A-to-I editing, their genomic sequence will be AT or/and AA respectively. To identify such cases, we have mapped available RNA-seq data from representative protostomian species (Methods). As the result, we did not find any non-canonical splice junctions in any of the species investigated (Table S3).

We have also used total RNA-seq data to annotate circular RNAs (circRNAs) in the tissues by detecting back-splicing events (*60*). As ADARs have been shown to antagonize back-splicing events by melting dsRNA structures within introns (*14*), we expected circRNAs in cephalopods to be not as abundant as in other animals. We have predicted circRNAs using total RNA-seq datasets by searching for backsplicing events (*60*), and rigorously filtered putative predictions to retain a set of high-quality 296 circRNAs (Methods). These RNAs were expressed in highly tissue-specific manner: the majority (200 / 296) were detected in only one of the tissues, and only 32 in more than 3 tissues.

#### Alternative cleavage and polyadenylation

We used FLAM-seq (*16*) to reconstruct mRNA 3'-untranslated regions (3'-UTRs). We found that the sequence context around cleavage sites in octopus strongly resembles that of other metazoans. More precisely, 71% of the sites were canonical metazoan polyadenylation signals (A(A/U)UAAA and variants) (Fig. S3 A-D). Thus, 3'-UTRs and their isoforms are likely generated by a canonical mechanism. The median length of 3'-UTRs in the octopus was approximately 380 nucleotides. As in other bilaterians, we find that genes expressed in the nervous system utilize, on average, longer 3'UTRs (Fig. S3 E) (*61*, *62*)**.** We also found 4,800 genes utilizing tandem alternative polyadenylation sites. As expected, distal cleavage sites usually contained stronger polyadenylation signals (Fig. S3 D) (*63*). Finally, we investigated poly(A) tail lengths. As in other animals, steady-state lengths of poly(A) tails are longer in neuronal tissues (*16*). Interestingly, poly(A) tails in the testis were shorter than in other tissues and contained a high proportion of guanosines (Fig. S3 F,G), which have not been described in any other animal.

Thus, while we found unusual poly(A) tails in the octopus testis, we conclude that the octopus employs polyadenylation patterns similar to those observed in other lophotrochozoans. In summary, the transcriptome of a common octopus does not show major departures from other lophotrochozoans in terms of alternative splicing diversity and rates, as well as in mRNA cleavage and polyadenylation. This matches a previous observation of gene content and intronic architecture of coleoids resembling that of other "slow-evolving" lophotrochozoans (*5*).

#### RNA editing index

A-to-I editing index has been computed as in (*64*). The main motivation for using this index instead of defining editing sites is the following. First of all, it avoids defining RNA editing sites explicitly using an arbitrary RNA sequencing depth threshold. Secondly, different genomic features have different coverage by the RNA sequencing reads. This uneven coverage leads to more editing sites being defined in the coding sequences per unit of length and may lead to false conclusion that coding sequences are edited at higher levels. Thus, when defining editing sites, the coverage should be accounted for. RNA editing index, on the other hand, is obtained by pulling the information from all reference adenosines for which there is any coverage present. This explicitly accounts for uneven coverage and thus allows comparison of different genomic regions. A hypothetical example illustrating this is shown in (Fig.S1A and B) When explicitly calling editing sites, feature A appears to be edited more (i.e. more editing sites per unit length) than feature B where no editing sites have been called due to low coverage. However, when comparing editing indices, the opposite is true. We have computed the editing index per each type of genomic feature in *O. sinensis* and *O. bimaculoides* genomes using the data generated in this study and (*5*) dataset respectively. Among the genic features in both species, the introns appear to have the highest editing indices followed by 3'-UTRs and coding sequences (Fig. S1).

#### Dependence between library sequencing depth and the number of captured miRNAs

To understand whether the sequencing depth in particular samples would affect the discovery of miRNAs in general, we compared the overall number of sequenced reads (raw) with the number of detected miRNAs in each sample (Fig. S5). For every sequencing library, we have recorded the number of captured "novel" miRNAs (i.e. miRNAs with evolutionary origins in Cephalopod lineage and younger) and the total number of all miRNAs. On average, 56.2% of miRNAs captured in neuronal tissues were novel compared to 43.2% of novel miRNAs in the non-neuronal tissues (p-value = 4.571e-05; Wilcoxon rank sum exact test). Even non-neuronal tissue with the highest coverage ("Suckers tip", 50 M reads; 53.6%) had the proportion of novel miRNAs lower than neuronal tissue with the lowest coverage ("Pedunculate and Olfactory lobe", 30 M reads; 57%). We thus conclude that the number of detected neuronal miRNAs is not elevated due to sample biases.

#### Editing of miRNAs

Changes in the seed sequence of a miRNA (positions 2-8 counted from the 5' end) are expected to alter target recognition (*26*). However, similar to mammals (*65*), our data show that miRNA seed regions are not edited to any considerable extent. In the mature miRNA sequences, the mismatches were more abundant towards the 3'-end of the molecule (Fig. S3A). The most common types of mismatches were non-templated 3' additions of cytosine, adenine and uridine (Fig. S7C). When pooling the data from all datasets, the average number of reads mapping to the position in a seed was 49,532 (median 27,967; minimum 12). 146/147 miRNAs have had all 7 bases of the seed profiled with at least 10 sequencing reads and 136 with at least 100 reads. The majority of mismatches was specific to either Truseq (6,443) or Clontech (2,355) datasets. In the remaining 894 cases, where a particular mismatch has been recovered by both library preparation methods, there was a mild correlation in the estimated mismatch frequencies (Spearman's rho = 0.46, correlation test p-value < 4.93E-43). In total, 5 miRNAs show signs of editing in their seed sequence above 1%. Of these, only 3 cases have been recovered in multiple tissues and none by both library preparation methods simultaneously (Table S4).

#### De-novo creation of miRNA targeting sites by RNA editing

Having established the higher conservation for miRNA response elements ("MRE's") compared to non-MRE controls (Main text, Methods), we tallied cases where A-to-I editing possibly created functional MREs by introducing A-to-G substitutions. We reasoned that if such "one-off MREs" (8-mers convertible to an MRE via A-to-G mismatch) exist and are functional, they would, similarly to canonical targeting sites, display higher conservation rates compared to other octamers (vis above). We recorded, genome-wide, the conservation of such one-off MREs and observed no trace of higher conservation when compared to G-to-A one-off controls (i.e. octamers convertible to an MRE via G-to-A substitution, Fig. 8E, p > 0.05, Wilcoxon rank sum test with continuity correction). Furthermore, we have checked whether one-off MREs with editing events had higher conservation rates compared to the same type of 8-mers not targeted by ADAR and found no difference. Out of 71,772 one-off MREs conserved between 2 octopus species (306 sequences), only 159 (57 different sequences) have been found to be targeted by ADAR. These 57 one-off MREs were conserved at the same rates as their unedited counterparts ($p > 0.05$; Wilcoxon rank sum test). This suggests that the de-novo creation of miRNA targeting sites via A-to-I editing may not be a widespread phenomenon in cephalopods. Similarly, A-to-I editing events with the potential to destroy the target sites have been found to happen rarely. Out of 10,053 MREs conserved between 2 octopus species, only 39 (0.3%) have been found to be targeted by ADAR (Methods). This is in line with the known preference of ADAR for the double-stranded RNA and the general depletion of the secondary RNA structure around functional miRNA targeting sites (*66*).

### Methods

#### Tissue dissection and RNA extraction

Adult specimens of O. vulgaris (body weight 800 g ± 50 g, mean ± SD) were collected from the Bay of Naples (Italy) and transferred to the Department of Biology, University of Naples Federico II (Italy). Our research was approved following the European Directive 2010/63 EU L276, the Italian DL. 4/03/2014, no. 26 and the ethical principles of Reduction, Refinement, and Replacement (Project n° 608/2016-PR-17/06/2016; protocol n°DGSAF 0022292-P-03/10/2017). Samples (N=8) were anesthetized by isoflurane insufflation (*67*), and tissues were dissected under sterile conditions following institutional guidelines. Tissue selected were: Axial nerve cords, Central heart, Vertical and Frontal lobes, Hepatopancreas, Ink sac, Intestine, Optic gland, Optic Lobe, Testes, Pedunculate lobe and Olfactory lobe, Posterior salivary glands, Retina, Skin, Stellate ganglion, Suckers at the base and the tip of the arm, Visceral (gastric) ganglion, White body.

Collected samples were snap frozen in liquid nitrogen and immediately put in Trizol, then stored at −80 °C. The RNA has been extracted from the tissues using Direct-zolTM RNA/ Miniprep (Zymo Research), following the manufacturer's protocol. The quality of RNA has been accessed with Bioanalyzer and only samples with intact RNA have been kept for library preparation.
A juvenile individual of *Octopus bimaculoides* (mantle size ~1 inch) was obtained from the National Resource Centre for Cephalopods (Galveston, Texas) on September 24$^{th}$, 2009. The specimen was shipped alive to Dartmouth College. The specimen was euthanised immediately by submersion directly into liquid nitrogen in a large mortar held in an ice bucket filled with dry ice. When completely frozen after a few minutes it was homogenised into a powder using a pestle. About 5 grams of homogenised powder was transferred to a 50 mL Oak Ridge screw cap centrifuge tube and mixed with Trizol for total RNA extraction using the standard protocol with glycogen added during the precipitation step (Invitrogen, Carlsbad).

Strings of eggs of *O. vulgaris* were obtained from the Instituto Español de Oceanografía (IEO, Tenerife, Spain). Embryos were incubated in a standalone system at KU Leuven, Belgium and collected at different developmental time points (St XI, XII.2, XIV, XVIII, one day old paralarvae). Embryos were dechorionated and the yolk was removed. Paralarval brains were dissected as described before (*38*). RNA was extracted from a pool of embryos or brains using Tri-reagent (Invitrogen) and the Qiagen Micro kit (Qiagen). All experiments involving hatchlings were approved by the ethical committee (permit P080/2021).

#### Poly-A+ mRNA library preparation and sequencing

Poly-A RNA sequencing libraries were prepared using the TruSeq Stranded mRNA Kit (Illumina) according to the manufacturer's instructions. Libraries were sequenced on a NextSeq 500 device at 1x76 cycles.

#### Total RNA library preparation and sequencing

300 ng of total RNA per sample was first depleted of ribosomal RNA using the RiboCop rRNA Depletion Kit (Lexogen, #144) according to the manufacturer's instructions. The rRNA-depleted

samples were then processed with the TruSeq mRNA stranded kit from Illumina. Libraries were then sequenced on a Nextseq 500 device at 2x76 cycles (paired-end).

#### Full-length mRNA library preparation and sequencing

Two full-length mRNA sequencing approaches used in this study have their own strengths and weaknesses and thus are complementary. In particular, FLAM-seq is biased towards shorter molecules, generating libraries of 1.5 Kbp median length. Iso-Seq, on the other hand, is susceptible to internal priming. These artifacts arise when oligo-dT primer aligns to an A-rich sequence inside mRNA instead of a poly(A) tail. The sequencing reads arising in the result appear truncated from a 3'-end and may be misinterpreted as alternative isoforms. FLAM-seq is insensitive to such an artifact as it replaces oligo dT priming with oligo dC priming onto an enzymatically added 3' guanosine/inosine anchor, and thus the sequenced mRNAs include the entire, non-templated poly(A) tail, which flags a detected mRNA 3' end as *bona fide*.In order to produce a comprehensive annotation of the *O. vulgaris* transcriptome, we therefore combined FLAM-seq with Iso-seq, with the addition of a size-selection step for enriching long transcripts which are underrepresented in FLAM-seq data. We reasoned that this strategy would combine the capacity of Iso-seq to generate extremely long cDNA molecules with the high accuracy of FLAM-seq in retrieving *bona fide* 3' ends of messenger RNA. For FLAM-seq library generation, the following detailed protocol was first applied to 4 micrograms of RNA from hepatopancreas: https://protocolexchange.researchsquare.com/article/pex-398/v1. On this sample, 16, 18, 20, or 22 PCR cycles were performed. The library profiles were checked on a bioanalyzer high-sensitivity DNA assay and concluded that 20 cycles yielded the best result in terms of library size and quantity. The complete protocol with 20 PCR cycles was therefore applied to 4 micrograms of the 6 samples listed in the Table S1, and the generated libraries were barcoded at the SMRTbell adapter ligation step and multiplexed on 6 Sequel SMRTcells in total.

For Iso-seq, the Iso-seq Express 2.0 workflow (PacBio) was applied to 500 nanograms of RNA from each of the 6 samples listed in the Table S1, using barcoded PCR primers for 14 cycles of PCR amplification, and performing size-selection after amplification with ProNex beads (Promega #NG2001), to enrich transcripts larger than 3 kb. Iso-seq libraries were then also multiplexed and sequenced on 6 Sequel SMRTcells in total. As indicated in the Table S1, an additional FLAM-seq library and Iso-seq library was generated in a second instance and again sequenced on 1 SMRTcell.

##### Processing of PacBio SMRT data

To obtain full-length non-chimeric reads (FLNCs) from Iso-Seq sequencing data, isoseq3 pipeline **(https://github.com/PacificBiosciences/IsoSeq)** has been used without a polishing step. We decided to skip a polishing step to increase the sensitivity for low-abundance transcripts. To obtain FLNC equivalents from FLAM-seq data, FLAMAnalysis pipeline has been used (*16*). In brief, the sequencing reads have been mapped to the genome using STARlong aligner (*68*)using default parameters. Next, poly(A) tails have been identified and trimmed from each read. We have used the resulting reads as an input for the next step.

#### Isoform reconstruction from FLNC reads

##### Read mapping to the genome and isoform reconstruction

For Iso-Seq, FLNC reads have been mapped to the *O. sinensis* genome using minimap2 (*69*):

```
 minimap2 -ax splice:hq -t {threads} -uf --secondary=no -C5 {params.genome} {input}
> {output}.
```

Next, the putative transcripts have been assembled for each tissue using TAMA Collapse (*70*):

```
python tama_collapse.py -s {input} -f {params.genome} -p
tama_collapse/{wildcards.sample} -x no_cap -e common_ends -c 99 -i 85 -a 10 -m 10 -
z 20 -sjt 10 -lde 5 -log {params.log}
```

FLAM-seq reads from FLAMAnalysis pipeline have been mapped to the genome with minimap2 using the same settings as above. As FLAM-seq provides precise resolution of the cleavage sites, TAMA Collapse has been used with a lower "three_prime_threshold" of 20 b.p. and a five prime threshold has been increased to 1000 b.p. effectively leading to a collapse of all reads mapping to the same cleavage site in the genome.

##### Merging isoforms from individual tissues

Isoforms from individual libraries have been merged with TAMA merge:

```
 tama_merge.py -m 10 -e common_ends -f {input.merge_file} -z 50 -a 10 -p
 {params.out_prefix} -d merge_dup
```

The merging resulted in 301,270 transcript models (86,394 if using previously polished models). In the merge file, higher weight has been given for FLAM-seq isoforms in determining 3'-end positions of the isoforms, and, conversely, higher weight has been given to Iso-Seq-derived isoforms in determining 5'-ends of the transcripts.

##### Isoform type classification and filtering

SQANTI2 tool (*71*) has been used to classify obtained isoforms with respect to the existing *O. sinensis* genome annotation:

```
 python {SQANTI2_PY} --skipORF --gtf --geneid --polyA_motif_list {input.polya_list} -t
 {threads} -n {params.chunks} -d {params.sqanti2
 _out} -e {input.exp} -c {input.splice} {input.isoforms_gtf} {GENOME_ANNO} {GENOME_FA}
```

The majority of isoforms supported by Isoseq data displayed elevated proportion of adenosines in the downstream genomic sequence thus suggesting their probable origin due to internal priming.
Putative models have been filtered using the following criteria:

1. All transcripts with putative reverse template switch artifacts (as defined by SQANTI2) have been filtered out
2. Full-splice match (FSM) transcripts have been retained only if the 3' end was reliable (<6 As in 10bp downstream genomic sequence)
3. Non-FSM transcripts have been retained only if:
    a. 3' end was reliable - either polyadenylation signal present or <6 As downstream

    b. Non-canonical junctions were supported by at least 5 uniquely mapping reads
    c. At least 2 reads were compatible with the isoform across all tissues
4. Fusion transcripts have been removed from the annotation entirely
5. Intergenic transcripts have been retained only if:
    a. There was an ORF>= 100AA predicted
    b. The model was supported by at least 5 reads
    c. The model was supported by conventional RNA-seq reads
    d. The model was multi-exonic
    e. 3' end was reliable (>6 As in 10bp downstream sequence)
    f. Passing splice junction support (vis above)
6. All antisense transcripts have been filtered
7. Finally, novel genes have been considered only if at least 1 of the associated transcripts is:
    a. Multi-exonic
    b. Splice junctions are supported (vis above)
    c. Supported by either FLAM-seq method only or both
    d. Supported by conventional RNA-seq

This filtering resulted in 59,579 mRNA isoforms associated with 10,957 reference genes. These isoforms have been added to original O. sinensis genome annotation (GCF_006345805.1_ASM634580v1). All the original isoforms with FSM isoforms from the newly predicted set have been removed. The genome annotation is available as Supplementary Data 1.

### Annotation of alternative splicing events

A Bioconductor SplicingGraphs (v 1.26.1; https://bioconductor.org/packages/release/bioc/html/SplicingGraphs.html) package has been used to count types of the alternative splicing events in the genome annotation obtained above by constructing and parsing splice graphs at each genomic locus as described in package vignette (https://bioconductor.org/packages/release/bioc/vignettes/SplicingGraphs/inst/doc/SplicingGraphs.pdf)

### Editing-generated splice sites

RNA-seq data from representative protostome animals (*O. bimaculoides* PRJNA285380, *N. pompilius* PRJNA614552, *C. gigas* PRJNA146329, *C. teleta* PRJNA379706) to corresponding genomes using STAR aligner (*68*) with the following parameters:

```
--alignIntronMin 20 --alignIntronMax 1000000  --alignSJoverhangMin 8
```

To make the datasets comparable, all sequencing reads have been trimmed to 50 bp and mapped to the respective genomes. The catalogs of splice sites provided by STAR aligner have been used to count the types of splice sites according to the intronic sequence.

### Comparison of alternative splicing rates across tissues

To compare alternative splicing rates across different tissues, we have used approach from previous comparative studies of alternative splicing (*50*, *51*)

**Exon skipping rates, rES**

Non-redundant exon triplets were extracted from the genome annotation .gff files using custom python scripts. For each such triplet, 3 junctions were generated by concatenating to 42 b.p. from the upstream and downstream exons:

- Exon1-exon2 (E1E2)
- Exon2-exon3 (E2E3)
- Exon2-exon3 (E1E3 an exon skipping event)

An effective mappability of each junction has been calculated by extracting 50-mers (max. 35) and mapping them back using bowtie to the set of all junctions. Only the triplets with all three junctions having mappability >= 20 have been kept for the downstream analysis. In the following analysis, the number of reads mapping to the junction has been adjusted by multiplying the read counts by 35/[effective mappability]. For the exon skipping rate analysis, only the reads not mapping to the genome have been used (i.e. those reads unmappable to the reference genome with bowtie). Exon skipping rate rES was then defined for each triplet as following:

$$\frac{N_{E1E3}}{(N_{E1E2}+N_{E2E3})/2+N_{E1E3}},$$

where $N_i$ represents mappability-adjusted number of reads mapping to a junction $i$.

**Intron retention rates, rIR**

Intron retention rates have been computed in a similar way but for the triplets consisting of the first bracketing exon (E1), intron (I) and the second bracketing exon (E2). Intron retention rate, rIR was then defined for each triplet as following:

$$\frac{(N_{E1I}+N_{IE2})/2}{(N_{E1I}+N_{IE2})/2+N_{E1E2}}$$

#### ES (IR) rate comparison across tissues

For every tissue, the triplets having at least 5 mapping reads to the major junctions (E1E2 + E2E2 for ES; E1I + IE2 for IR) and a minimal mappability of all 3 junctions have been retained. Then, 1000 triplets from this set have been chosen randomly and exactly 25,000 reads have been mapped to the junctions. Then, the rES (rIR) rate were determined as described above. This approach has been repeated 100 times for every tissue in the dataset.

#### Annotation of circular RNAs

FindCirc2 ([https://github.com/rajewsky-lab/find_circ2](https://github.com/rajewsky-lab/find_circ2)) (*60*) has been used to annotate circular RNAs from the total RNA-seq data. Putative circRNA predictions have been rigorously filtered to ensure high evidence of the backsplice junctions without any other explanations for the read mapping (such as trans-splicing events):
WARN_OTHER_CHROM_MATE==0 and WARN_OUTSIDE_SPLICE_JUNCTION==0 and SUPPORT_INSIDE_MATE>=5 and WARN_OUTSIDE_MATE<5 and WARN_OUTSIDE_SPLICE_JUNCTION==0 and WARN_UNRESOLVED_EXTRA_BACKSPLICE==0 and WARN_UNRESOLVED_LINSPLICE==0

Finally, only back-splice junctions corresponding to known splice donor and acceptor sites have been kept.

### Alternative cleavage and polyadenylation

#### 3’-UTR annotation with FLAM-seq

To annotate 3’-UTRs in the octopus genome, we have used FLAM-seq data as this method avoids artifacts caused by internal priming. Each FLAM-seq read comes from an individual mRNA molecule (FLAM-seq uses unique molecule identifiers) and contains its full sequence as well as the sequence of the poly(A) tail. First, the reads were processed by a FLAMAnalysis pipeline (https://github.com/rajewsky-lab/FLAMAnalysis) to estimate the start position of a poly(A) tail and trim it from the sequence. Next, the resulting reads have been mapped to the genome using minimap2 (v 2.17-r941) using the parameters "--cs -ax splice:hq -uf --secondary=no -C5", and the 3'-most position of each read has been recorded generating ~110,000 initial putative cleavage sites. The tags closer that 20 base pairs have been merged, and their counts have been assigned to the most implicated site in the cluster. Then, the clusters were merged if separated by less than 40 nucleotides (Fig. S3A). Some genes may be missing from a genome annotation. To prevent sampling from such genes which would induce misassignment of the FLAM-seq tags, we kept putative cleavage sites only if:

1. There is a continuous short RNA-seq coverage of at least 5 reads from the FLAM-seq tag to a stop codon of an upstream gene, or
2. At least one read assigned to the cluster overlaps the stop codon of the gene the cluster is assigned to.

**40,949** putative cleavage sites have passed the filtering above and were included in the final dataset. This final set of cleavage sites contains 16,573 FLAM-seq clusters assigned to 7593 genes (Fig. S3A).

#### Cleavage sites and polyadenylation signals

##### Sequence profiles around cleavage sites

The sequence composition around cleavage sites for *D. melanogaster* and *C. elegans* (fig. S2A) were obtained by extracting the regions around 3’-ends of gene models (both *D. melanogaster* and *C. elegans* genome annotations were obtained from EnsemlMetazoa (*C. elegans* genome assembly WBcel235; *D. melanogaster* genome assembly BDGP6.32).

##### Polyadenylation signals

To determine polyadenylation signals (PASs) in the 3’UTRs, we queried the sequence upstream of the cleavage sites (50 nt) for a presence of polyadenylation signals described in other metazoans. In cases of multiple PASs occurring in the sequence, the one that’s more abundant in the whole dataset has been selected.

##### 3’-UTR and poly(A) length and composition in the tissues

In the dataset, genes with higher coverage would theoretically exhibit longer maximal 3’-UTRs as there would be a higher chance of capturing those. However, the relationship between the sequencing depth per gene and the maximal recovered 3’-UTR length is weak and explained about 1% of the variability in the 3’-UTR length (data not shown). We thus concluded that our FLAM-seq dataset allows for a robust 3’-UTR length estimation for sampled genes. For each gene, a mean distance from the FLAM-seq tags to the annotated stop codon has been computed across all tissues in the dataset. Different genes showed profound differences in 3’-UTR lengths based on whether they have multiple cleavage sites annotated and their tissue expression patterns. Genes with multiple annotated 3’-UTRs exhibited longer 3’-UTRs compared to genes with only one cleavage site captured. The genes specifically expressed in the nervous tissues utilized the longest 3’-UTRs (fig S3). In addition to the lengths of 3’-UTRs, we have investigated steady-state lengths of mRNA poly(A) tails. Similar to the 3’-UTRs, poly(A) tails were longer in the neuronal tissues (123 - 138.5 nt) than in non-neuronal (45-110 nt). The shortest poly(A) tails have been observed in the testis (median 45.0 nt). In addition, mRNA tails in testis showed unusually high percentage of guanosines (up to 10%) (Fig. S3)

#### A-to-I editing analyses

##### A-to-I editing detection

To call editing events, short-read RNA-seq data generated in this study were mapped to the genome with STAR aligner (*68*). Alignment files were filtered to retain only uniquely mapping reads. The mismatches between the genome assembly and the alignments were called with bcftools (*72*). The same approach was used for DNA-seq data from an *O. vulgaris* genome assembly project (*55*). Variants occurring in RNA-seq data but not in DNA-seq have been retrieved with bcftools isec. All genomic loci with more than 10 mapping reads of which at least 3 contain guanosine instead of adenosine were considered editing sites. This approach led to identification of 68,338 putative editing sites.

##### A-to-I editing index

Briefly, for every genomic feature, the total numbers of adenosines and guanosines sequenced at all positions with reference A’s were determined. Then, an editing index was computed as the number of guanosines divided by a total number of sequenced bases.

#### Small RNA sequencing and analysis

##### Small RNA library preparation and sequencing

**Octopus vulgaris**
Small RNA sequencing libraries were prepared using 2 different kits: the 1st one is SMARTer smRNA-Seq kit for Illumina from Clontech according to manufacturer’s instruction using 10ng total RNA, the libraries were pooled together with 20% phix and sequenced on the NextSeq 500, 1x51. The 2nd Kit is the Truseq Small RNA Kit from (Illumina), the libraries were prepared according to manufacturer’s

instruction using between 100 ng and 1000ng Total RNA the libraries were pooled together and sequenced on the NextSeq 500, 1x51

**Octopus bimaculoides**
SmallRNA libraries prepared at the Yale University School of Medicine W. M. Keck facility using standard manufacturers' protocol and sequenced on the Illumina Genome Analyzer II platform loaded onto a single lane.

**Octopus vulgaris (developmental stages)**
The sequencing libraries have been prepared using SMARTer smRNA-Seq kit for Illumina from Clontech according to manufacturer's instructions.

**Quality control and expression quantification**
The quality control of the sequencing data has been performed using miRTrace software (v 1.0.1, https://github.com/friedlanderlab/mirtrace) (*73*). MirTrace QC files are available as a part of supplementary data. MiRDeep2 tool v2.0.1.2 (*74*) was used to quantify miRNA expression in the tissues.

##### miRNA Annotation

Generally, we followed the annotation procedures described by (*75*). For conserved miRNA discovery we used MirMachine (https://github.com/sinanugur/MirMachine). Briefly, covariance models for each conserved miRNA family in MirGeneDB (*46*) were searched in the genomes of *Octopus vulgaris* and *O. bimaculoides,* the bobtail squid *Euprymna scolopes* and *Nautilus pompilius*. Hits for Eumetazoan, Bilaterian, Protostomian, Lophotrochozoan and Molluscan families/genes were returned, respectively. After this, for Octopus species, and squid (but not the Nautlius) species specific and quality-filtered miRNA sequencing datasets were pooled and used in MirMiner (*76*) for novel miRNA gene discovery in each species. Conserved and novel predictions were compared across cephalopods and other mollusc species in MirGeneDB. Annotation of miRNA families, genes and paralogues was conducted using synteny and orthology information combined with sequence comparison (with emphasis on seed regions).

#### miRNA editing analysis

To detect A-to-I editing events in the miRNAs, small RNA sequencing data has been mapped to predicted mature sequences with bowtie allowing maximum 2 mismatches and no multi-mapping reads. Resulting alignment (.bam) files have been processed with a custom python script to return base counts at every position of the reference. Then, for the reference positions with adenosines having sufficient coverage (more than 10 reads), the proportion of guanosines has been computed. For a Figure S8, the maximum such proportion in miRNA seed ( position $2^{nd}$-$8^{th}$) has been plotted with ComplexHeatmap R package. (*77*)

### miRNA response-element conservation analysis

#### Alignment of orthologous 3'-UTRs

**11,361** one-to-one ortholog pairs between *O. sinensis* and *O. bimaculoides* have been identified using OrthoFinder2 (*78*) with default parameters and proteomes of 17 other representative metazoans (*Aplysia californica*, *Biomphalaria glabrata, Branchiostoma floridae, Capitella teleta, Ciona intestinalis, Crassostrea gigas, Euprymna scolopes, Homo sapiens, Lingula anatina, Lottia gigantea, Mus musculus, Nautilus pompilius, Nematostella vectensis, Octopus minor, Pinctada fucata, Saccoglossus kowalewskii, Strongylocentrotus purpuratus*). Then, the coding sequences from *O. sinensis* have been aligned to the genome of *O. bimaculoides* using GMAP (--cross-species, --min-identity=0.6) (*79*). The alignments have been filtered to include only the cases, where the best alignment for a transcript was the alignment to a corresponding ortholog and where the end of CDS has been aligned *precisely* to the stop codon of the reference genome. This filtering has resulted in **7,969** alignments. Next, the genomic sequence downstream of the alignment has been extracted. Finally, each pair of sequences corresponding to the orthologous 3'UTRs has been aligned with Clustal Omega (*80*) using default parameters.

#### K-mer conservation scores

The "conservation score" between 2 *Octopus* species has been then obtained by computing the fraction of cases where a k-mer in reference (*O. sinensis*) is matched exactly by the k-mer in the query (*O. bimaculoides*) for every distinct k-mer. For the analysis, we kept only k-mers present at least 10 times in the alignments. We considered miRNA and mRNA "co-detected" if they have been recovered at 100 and 10 counts-per-million respectively in any of the tissues in the dataset.

#### One-off MRE conservation

The conservation rates of 8-mers potentially convertible to MREs via a single editing event (one-off MREs) were compared to the conservation rates of control 8-mers with a similar dinucleotide composition obtained by reshuffling as well as 8-mers convertible to MREs via G-to-A substitution.

#### Editing of MREs and one-off MREs

To determine whether ADAR is targeting putative MREs and one-off MREs, all such 8-mers conserved between 2 octopus species have been intersected with A-to-I editing events identified previously. To ensure that the absence of observed editing is not a result of a missing data, only 8-mers with a total RNA-seq coverage of more than 10 mapping reads have been considered.

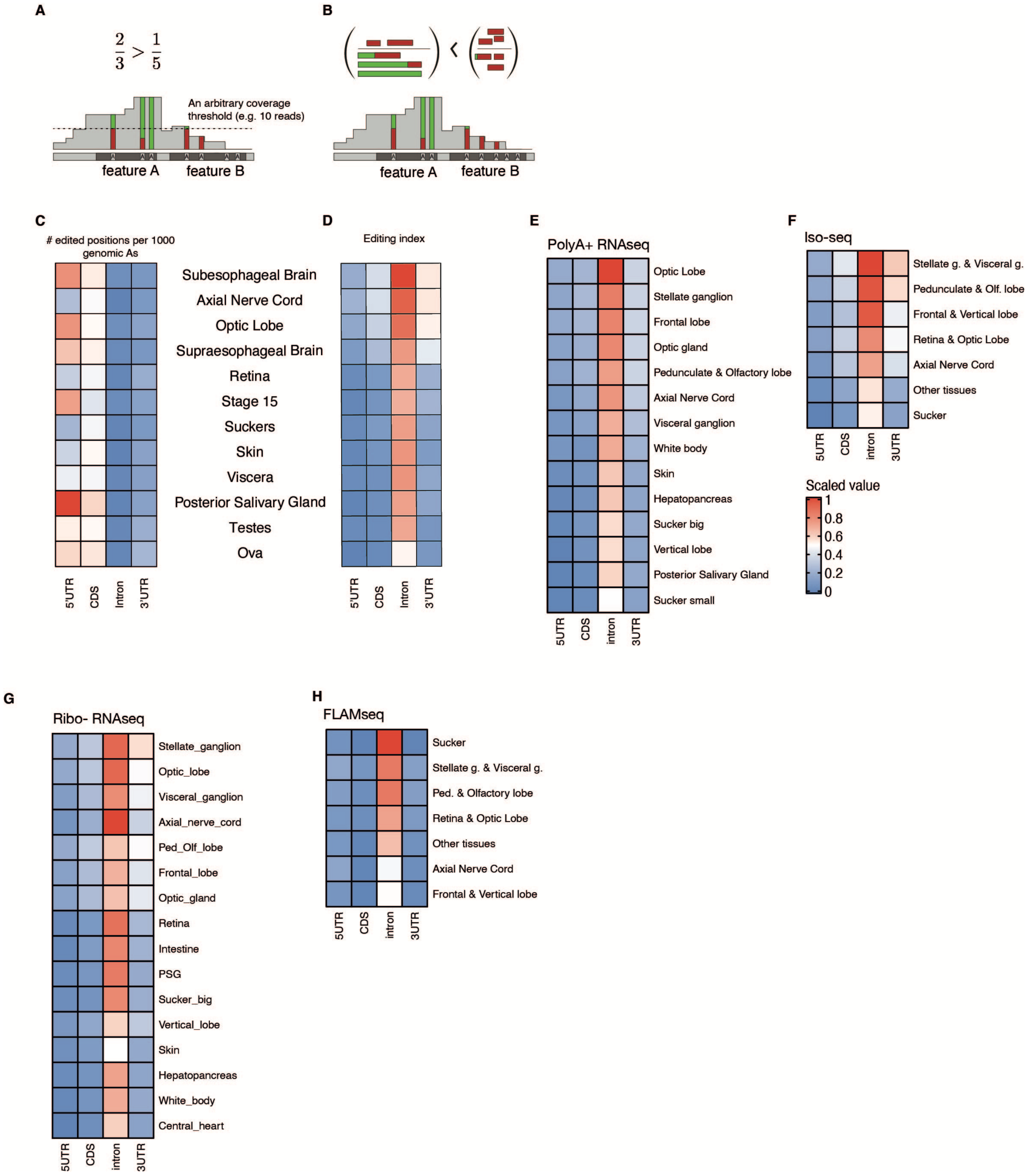


**Fig. S1. Genomic targets of A-to-I editing**

**(A)** Definition of calling editing sites. Editing sites are required to have minimal coverage (e.g., 10 RNA-seq reads) to increase the precision. Some genomic features (e.g., introns) will not have coverage above the threshold across their whole length. Thus, normalization by feature-length will lead to the underestimation of editing intensity in such loci.
**(B)** Editing index approach introduced in (*64*). A-to-G mismatches are pooled across the whole feature. Note that the index implicitly accounts for the coverage and uncovered positions have no effect on the resulting value.
**(C** and **D)** An average number of called editing sites (C) or editing index (D) per type of genomic feature in each tissue of *O. bimaculoides* (*5*). The values have been normalized such that the maximal and minimal value per matrix is 1 and 0 respectively. While neuronal tissues show higher values in both measures, the most edited features differ. e-h.
**(E-G)** Average editing indices of genomic features of *O. vulgaris* for every RNA-sequencing dataset generated in this study.
**(I)** The difference in editing intensity between DNA-binding domains and the remaining portion of the protein for identified transcription factors (for details vis Supplementary Information). Domain types have been sorted according to their frequency; all domains with only one protein have been merged into "Others".

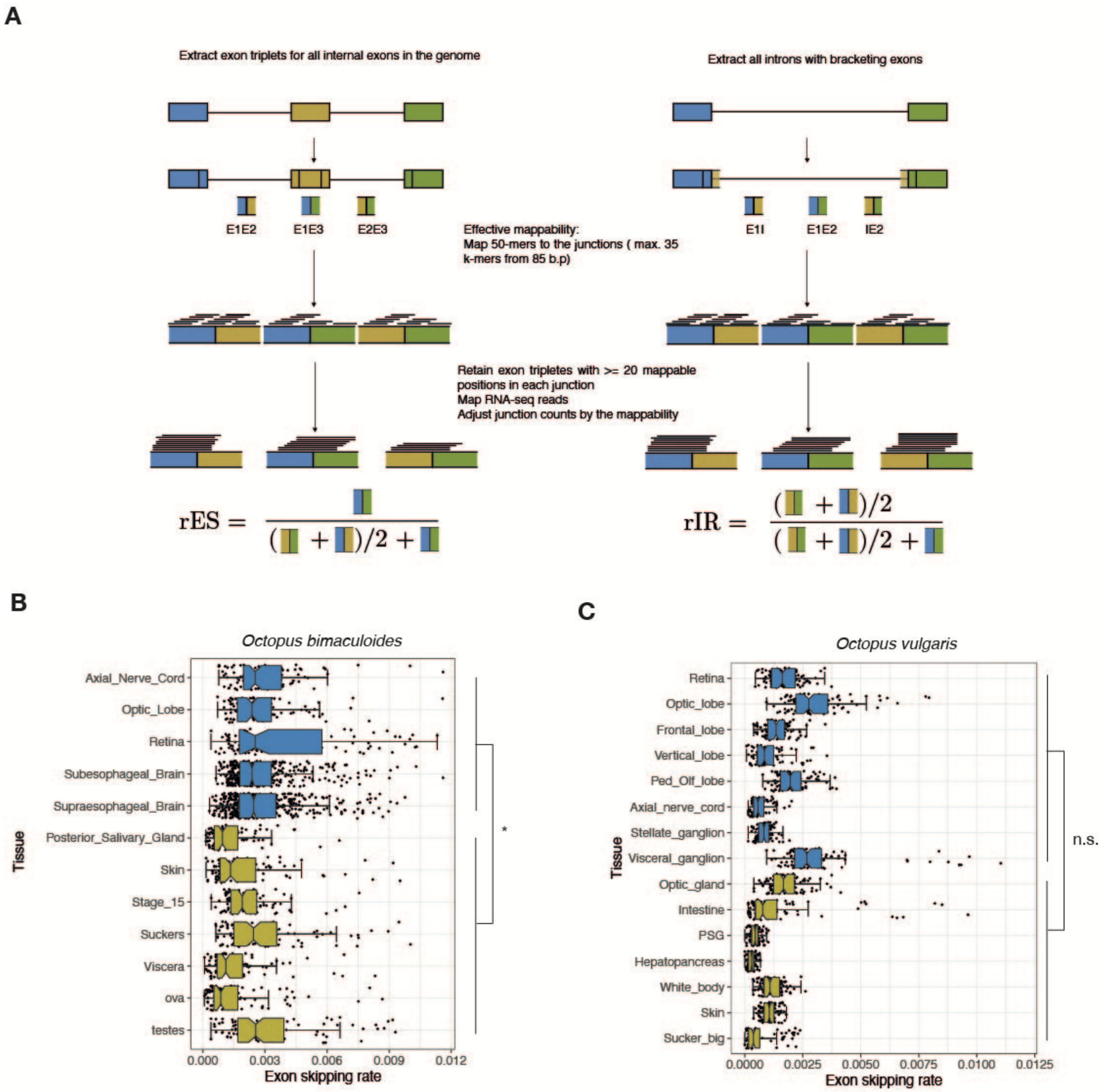


**Fig. S2. Alternative splicing rates across tissues**

**(A)** General approach used for quantification exon skipping and intron retention rates; adapted from (*50*). **(B** and **C)** Exon skipping rates in representative tissues of *O. bimaculoides* (**B**) and *O. vulgaris* (**C**) (Methods). Wilcoxon rank sum test with continuity correction was used to test for the higher median per-tissue exon skipping rate between neuronal and non-neuronal tissue types, * $p < 0.05$, ** $p < 0.01$, *** $p < 0.001$

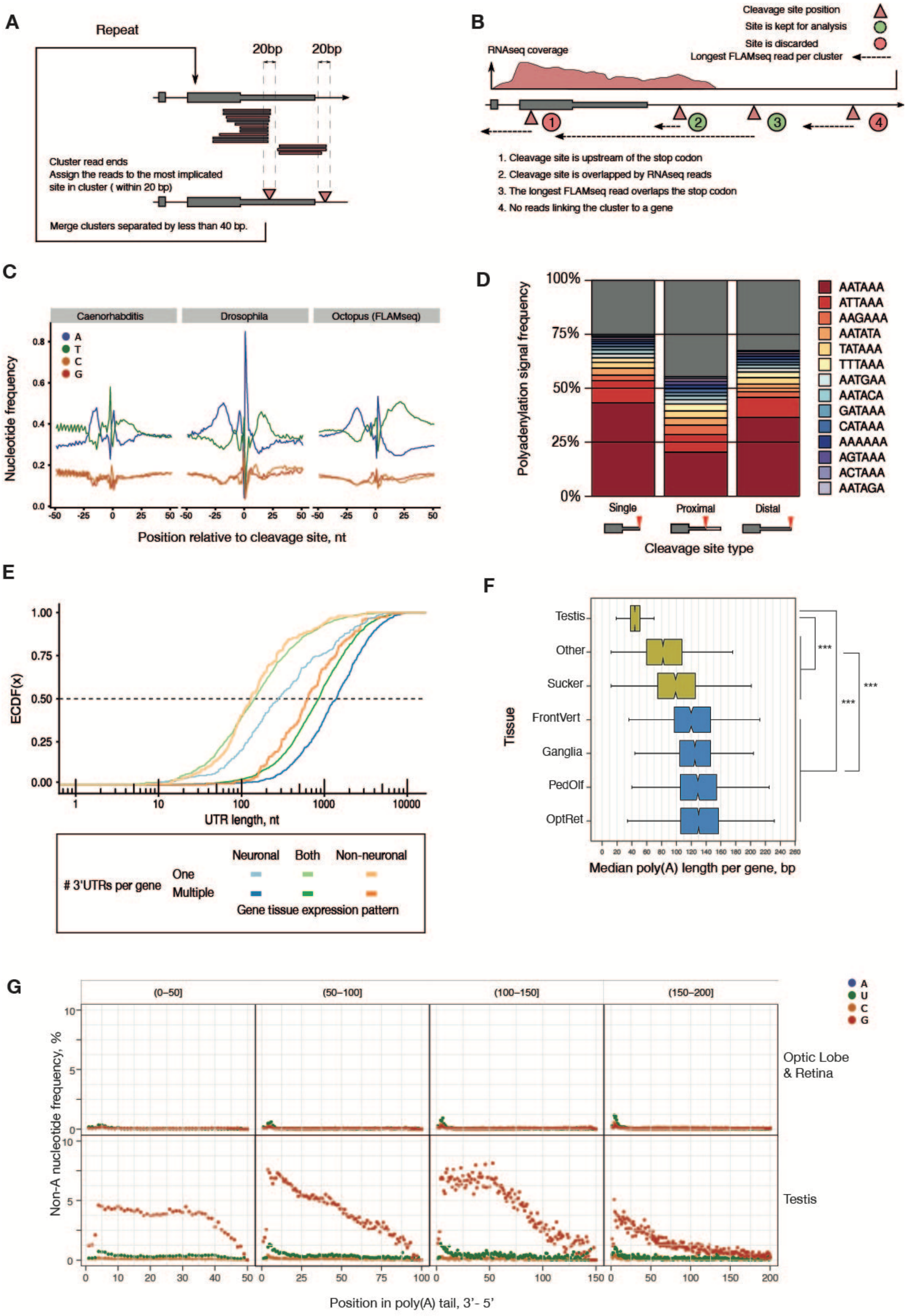
A
Repeat
20bp
20bp
Cluster read ends
Assign the reads to the most implicated site in cluster ( within 20 bp)
Merge clusters separated by less than 40 bp.
B
Cleavage site position
Site is kept for analysis
Site is discarded
Longest FLAMseq read per cluster
RNAseq coverage
1. Cleavage site is upstream of the stop codon
2. Cleavage site is overlapped by RNAseq reads
3. The longest FLAMseq read overlaps the stop codon
4. No reads linking the cluster to a gene
C
Caenorhabditis
Drosophila
Octopus (FLAMseq)
Nucleotide frequency
Position relative to cleavage site, nt
D
Polyadenylation signal frequency
AATAAA
ATTAAA
AAGAAA
AATATA
TATAAA
TTTAAA
AATGAA
AATACA
GATAAA
CATAAA
AAAAAA
AGTAAA
ACTAAA
AATAGA
Single
Proximal
Distal
Cleavage site type
E
ECDF(x)
UTR length, nt
### 3'UTRs per gene
One
Multiple
Neuronal
Both
Non-neuronal
Gene tissue expression pattern
F
Testis
Other
Sucker
FrontVert
Ganglia
PedOlf
OptRet
Tissue
Median poly(A) length per gene, bp
G
(0–50]
(50–100]
(100–150]
(150–200]
Optic Lobe & Retina
Testis
Non-A nucleotide frequency, %
Position in poly(A) tail, 3'- 5'

**Fig. S3. mRNA cleavage and polyadenylation**

**(A)** A general approach of assigning FLAM-seq tags to genes.
**(B)** A hypothetical locus with examples of putative mRNA cleavage sites with various supporting evidence and different results of filtering
**(C)** Sequence profiles around cleavage sites in representative protostomian species.
**(D)** Proportions of polyadenylation signals identified within 40 b.p upstream of the cleavage sites.
**(E)** Empirical cumulative distribution function of 3'-UTR length for various groups of genes in the *O. sinensis* genome.
**(F)** Median mRNA poly(A) tail lengths per gene in different tissues as determined by FLAMseq. Only genes with at least 5 distinct UMIs are shown. (Supplementary Text). Wilcoxon rank sum test with continuity correction, * $p< 0.05$,** $p < 0.01$, *** $p < 0.001$
**(G)** Non-adenosine nucleotide proportions across mRNA tails in testis and neuronal tissue (optic lobe and retina). The tails have been grouped into 4 length bins and aligned such that 0 refers to the 3'-most base.

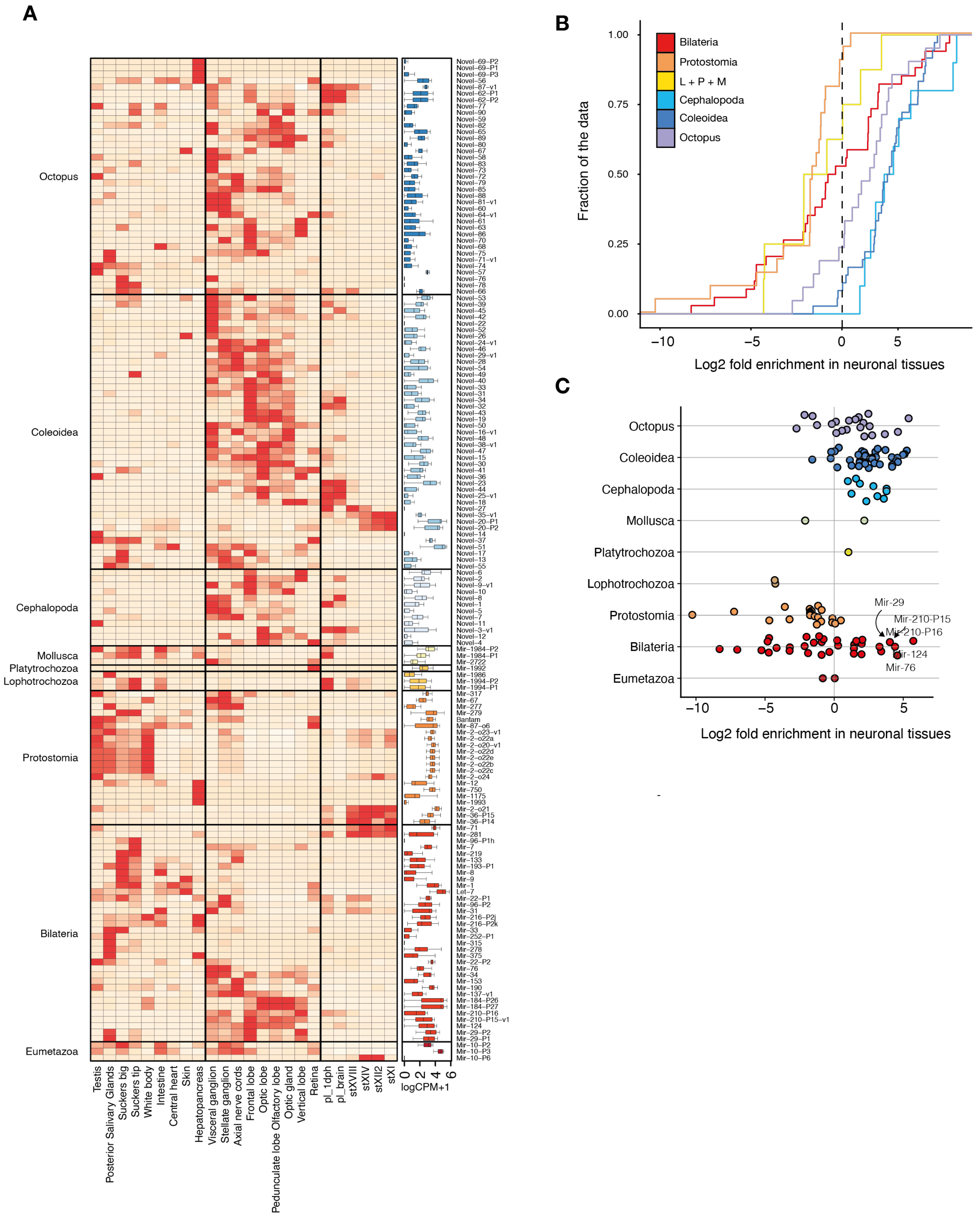
A
Octopus
Coleoidea
Cephalopoda
Mollusca
Platytrochozoa
Lophotrochozoa
Protostomia
Bilateria
Eumetazoa
Testis
Posterior Salivary Glands
Suckers big
Suckers tip
White body
Intestine
Central heart
Skin
Hepatopancreas
Visceral ganglion
Stellate ganglion
Axial nerve cords
Frontal lobe
Optic lobe
Peduncluate lobe Olfactory lobe
Optic gland
Vertical lobe
Retina
pl_1dph
pl_brain
stXVIII
stXIV
stXII2
stXI
0 2 4 6
logCPM+1
B
Bilateria
Protostomia
L + P + M
Cephalopoda
Coleoidea
Octopus
Fraction of the data
1.00
0.75
0.50
0.25
0.00
−10
−5
0
5
Log2 fold enrichment in neuronal tissues
C
Octopus
Coleoidea
Cephalopoda
Mollusca
Platytrochozoa
Lophotrochozoa
Protostomia
Bilateria
Eumetazoa
Mir-29
Mir-210-P15
Mir-210-P16
Mir-124
Mir-76
−10
−5
0
5
Log2 fold enrichment in neuronal tissues

**Fig. S4. Extended expression patterns and A-to-I editing of miRNAs**

**(A)** Detailed miRNA expression heatmap as in Fig.1. Boxplots on the right summarize distributions of counts-per-million (CPMs) across tissues for every miRNA.
**(B** and **C)** Log2 fold enrichment in neuronal tissues. Only miRNAs detected with at least 3 CPMs in both neuronal and non-neuronal tissues are shown. Enrichment is defined as the ratio of average expression values in neuronal and non-neuronal tissue types. In (**B)**, miRNAs with origins in Lophotrochozoa, Platytrochozoa, and Mollusca lineages have been grouped together (L + P + M) due to a low number of genes in individual groups; for the same reason, miRNAs of Eumetazoan origin have been omitted. In **C**, top 5 of the bilaterian miRNAs with the highest neuronal enrichment are labelled.

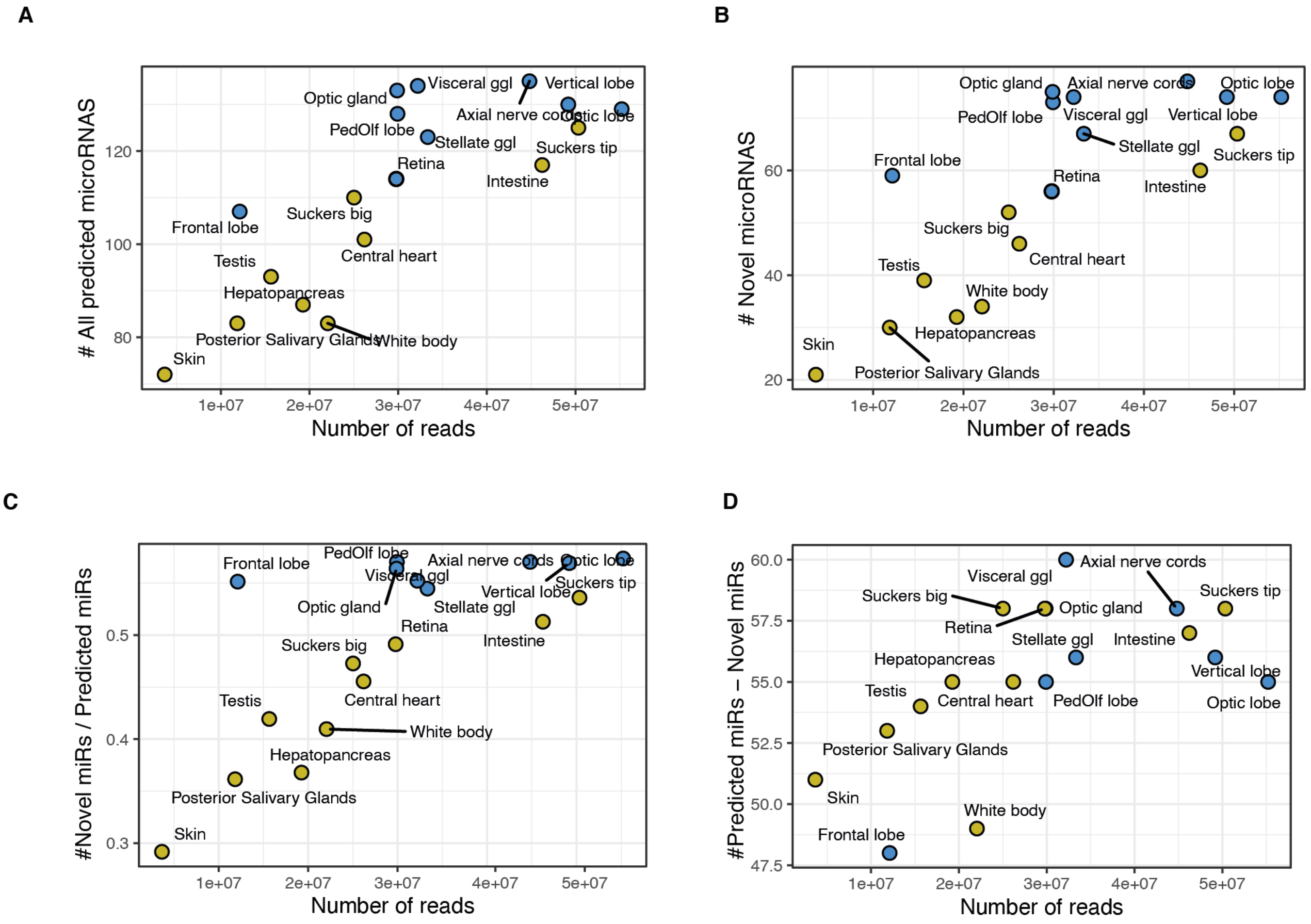


**Fig. S5. Sequencing depth and miRNA discovery**

**(A)** Relationship between sequencing depth (x-axis) per library and the number of captured miRNAs. Panel (**B)** depicts the numbers of "novel" miRNAs (i.e. with origin in Cephalopoda lineage and younger) while panel **(C)** reports relative proportions of novel miRNAs among all captured miRNAs. **(D)** Number of evolutionary "old" miRNAs detected in every sequencing library.

**A**

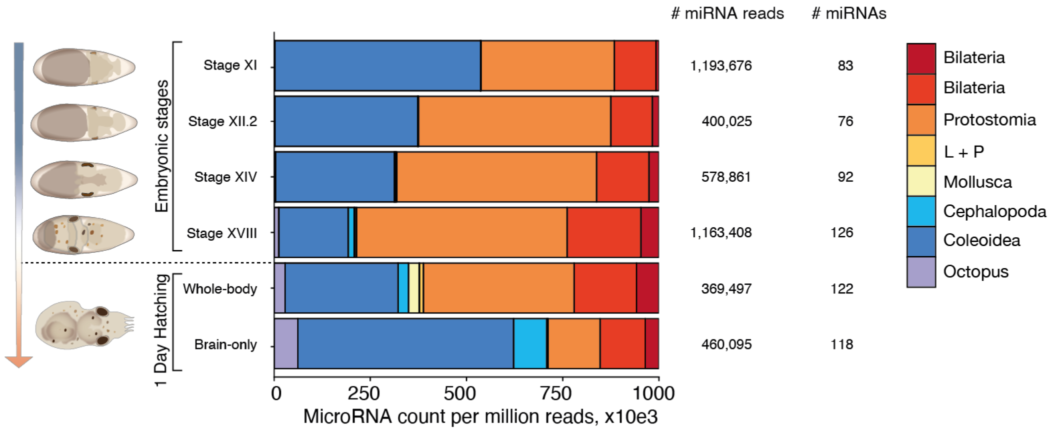

### miRNA reads
### miRNAs
Embryonic stages
1 Day Hatching
Stage XI
Stage XII.2
Stage XIV
Stage XVIII
Whole-body
Brain-only
1,193,676
83
400,025
76
578,861
92
1,163,408
126
369,497
122
460,095
118
Bilateria
Bilateria
Protostomia
L + P
Mollusca
Cephalopoda
Coleoidea
Octopus
0
250
500
750
1000
MicroRNA count per million reads, x10e3


**B**

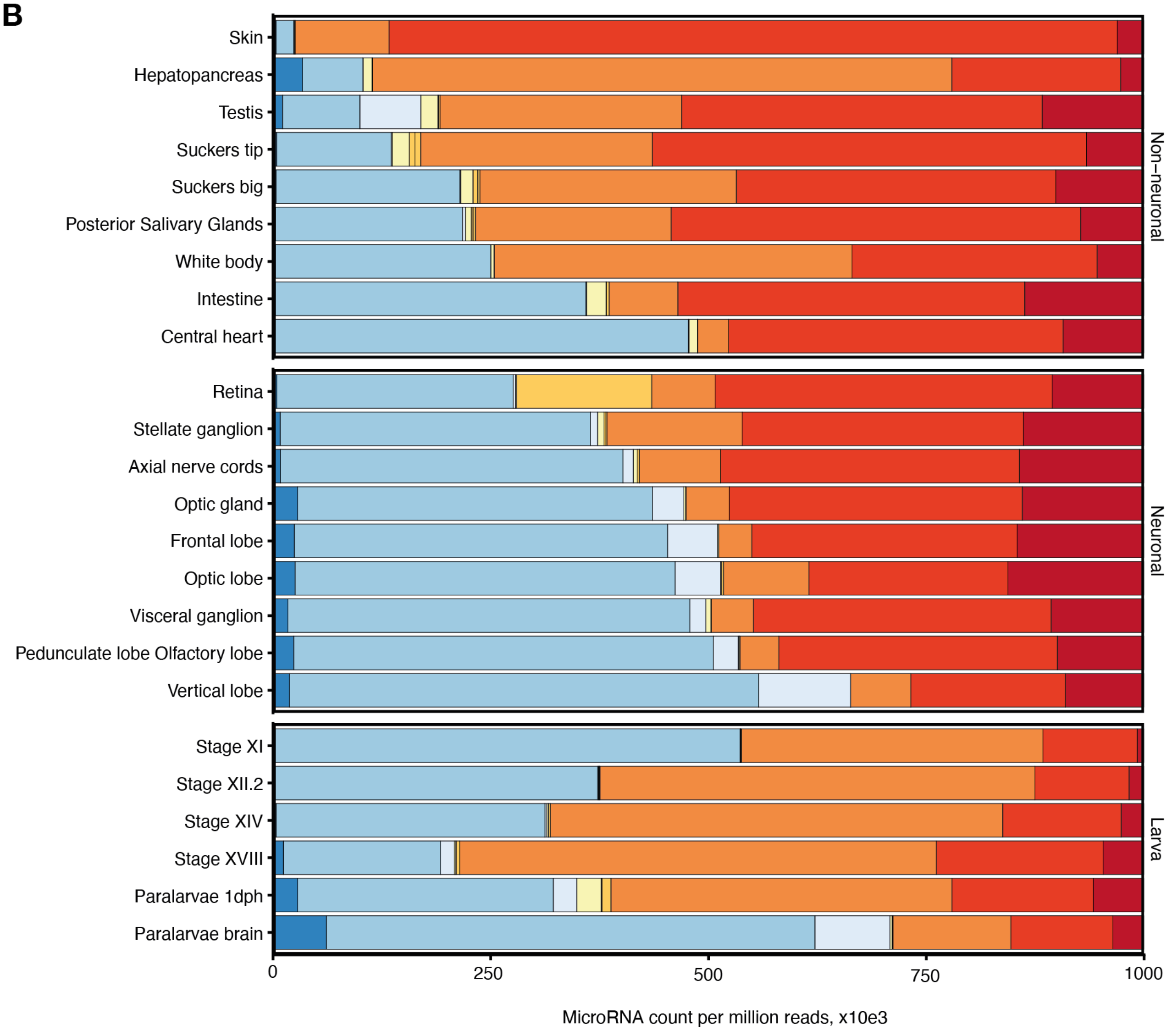

Skin
Hepatopancreas
Testis
Suckers tip
Suckers big
Posterior Salivary Glands
White body
Intestine
Central heart
Non–neuronal
Retina
Stellate ganglion
Axial nerve cords
Optic gland
Frontal lobe
Optic lobe
Visceral ganglion
Pedunculate lobe Olfactory lobe
Vertical lobe
Neuronal
Stage XI
Stage XII.2
Stage XIV
Stage XVIII
Paralarvae 1dph
Paralarvae brain
Larva
0
250
500
750
1000
MicroRNA count per million reads, x10e3

**Fig. S6. Fraction of transcriptome dedicated miRNAs of different phylogenetic ages**

**(A)** An expanded version of Fig. 4 with the total numbers of miRNA-derived reads and the numbers of miRNAs detected per library.
**(B)** MiRNA expression as reads per million colored according to the phylogenetic node of origin in all tissues profiled in the study. Non-neuronal and neuronal tissues have been sorted according to the proportion of reads assigned to coleoid miRNAs.

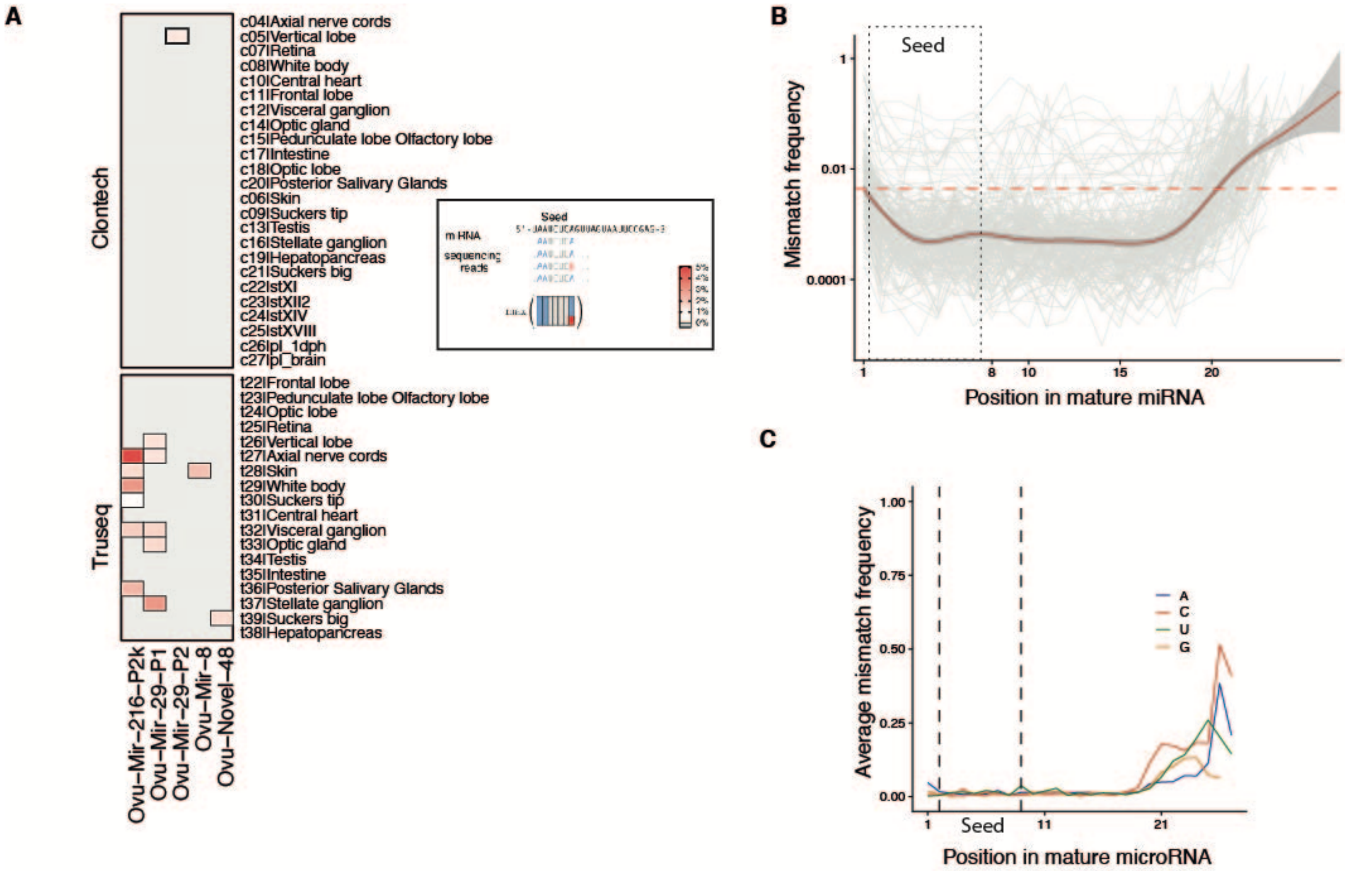


**Fig. S7 Editing of miRNAs**

**(A)**. Heatmap of maximal editing levels captured within miRNA seeds for all 5 miRNAs with editing levels above 1% in at least one of the tissues.

**(B)**. Total proportion of mismatches recovered for every position in individual miRNAs (gray lines). The horizontal red line marks a median sequencing error rate for Illumina NextSeq machine used in the study ( 0.429% from (*81*)).

**(C)**. Total proportion of mismatches called at different positions of miRNAs separated by the alternative base type.

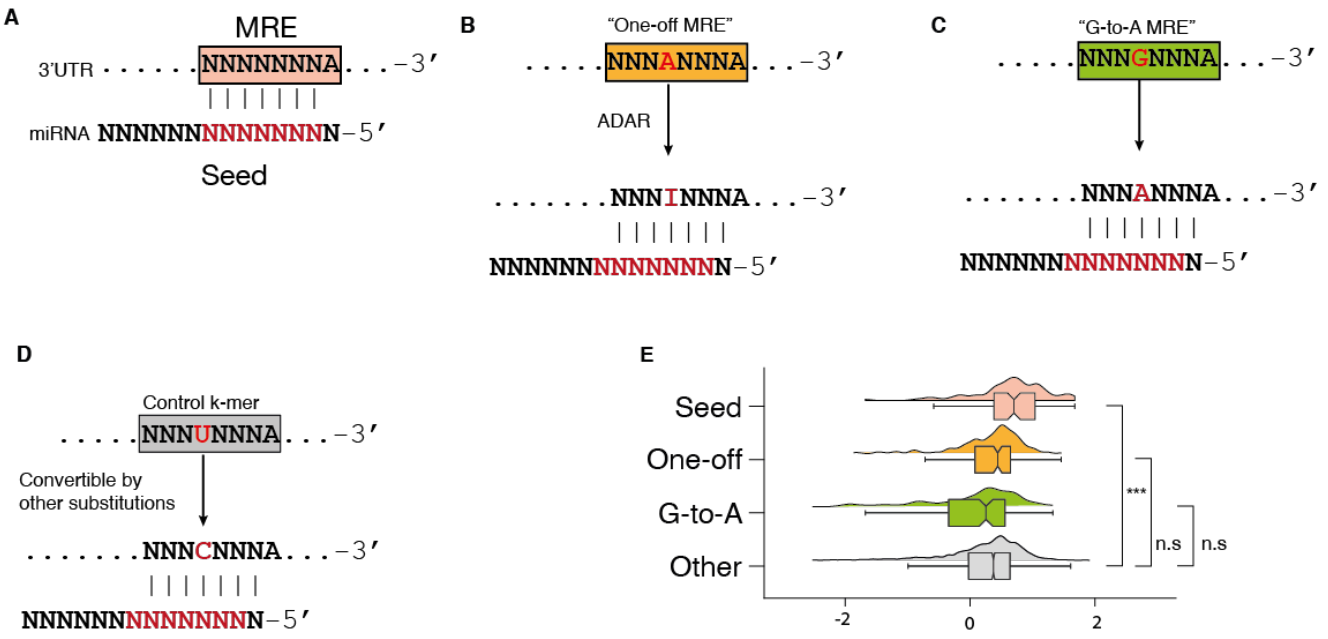


**Fig. S8 Conservation of 8-mers potentially convertible to MRE by A-to-I editing**

**(A** to **D)** Diagrams illustrating the types of k-mers compared in the panel **(E)**. "One-off MRE" – an 8-mer potentially convertible to MRE via A-to-I substitution; "G-to-A MRE" – an 8-mer convertible to MRE by a single G-to-A substitution; "Other" – 8-mers differing by 1 base from MREs and thus convertible to MRE via a single substitution that is not A-to-I or G-to-A. **(E)** Conservation Z-scores for the 8-mers (Supplementary Text; Wilcoxon rank sum test with continuity correction, * $p< 0.05$,** $p < 0.01$, *** $p < 0.001$)
